# Criterion Validity and Relationships between Alternative Hierarchical Dimensional Models of General and Specific Psychopathology

**DOI:** 10.1101/2020.04.27.064303

**Authors:** Tyler M. Moore, Antonia N. Kaczkurkin, E. Leighton Durham, Hee Jung Jeong, Malerie G. McDowell, Randolph M. Dupont, Brooks Applegate, Jennifer L. Tackett, Carlos Cardenas-Iniguez, Omid Kardan, Gaby N. Akcelik, Andrew J. Stier, Monica D. Rosenberg, Donald Hedeker, Marc G. Berman, Benjamin B. Lahey

## Abstract

Psychopathology can be viewed as a hierarchy of correlated dimensions. Many studies have supported this conceptualization, but they have used alternative statistical models with differing interpretations. In bifactor models, every symptom loads on both the general factor and one specific factor (e.g., internalizing), which partitions the total explained variance in each symptom between these orthogonal factors. In second-order models, symptoms load on one of several correlated lower-order factors. These lower-order factors load on a second-order general factor, which is defined by the variance shared by the lower-order factors. Thus, the factors in second-order models are not orthogonal. Choosing between these valid statistical models depends on the hypothesis being tested. Because bifactor models define orthogonal phenotypes with distinct sources of variance, they are optimal for studies of shared and unique associations of the dimensions of psychopathology with external variables putatively relevant to etiology and mechanisms. Concerns have been raised, however, about the reliability of the orthogonal specific factors in bifactor models. We evaluated this concern using parent symptom ratings of 9-10 year olds in the ABCD Study. Psychometric indices indicated that all factors in both bifactor and second-order models exhibited at least adequate construct reliability and estimated replicability. The factors defined in bifactor and second-order models were highly to moderately correlated across models, but have different interpretations. All factors in both models demonstrated significant associations with external criterion variables of theoretical and clinical importance, but the interpretation of such associations in second-order models was ambiguous due to shared variance among factors.

**General Scientific Summary:** Some investigators have proposed that viewing the correlated symptoms of psychopathology as a hierarchy in which all symptoms are related to both a general (p) factor of psychopathology and a more specific factor will make it easier to distinguish potential risk factors and mechanisms that are nonspecifically related to all forms of psychopathology versus those that are associated with specific dimensions of psychopathology. Parent ratings of child psychopathology items from the Adolescent Brain Cognitive Development (ABCD) Study were analyzed using two alternative statistical models of the proposed hierarchy. All factors of psychopathology defined in both bifactor and second-order models demonstrated adequate psychometric properties and criterion validity, but associations of psychopathology factors with external variables were more easily interpreted in bifactor than in second-order models.

---

Many scholars argue that psychopathology should not be conceptualized as distinct categorical diagnoses, but as a hierarchy of correlated dimensions of maladaptive behaviors, emotions, and cognitions (Achenbach, Conners, Quay, Verhulst, & Howell, 1989; Helzer, Kraemer, & Krueger, 2006; Krueger et al., 2018; Markon, Chmielewski, & Miller, 2011). Recently, this hierarchy has been posited to consist of a broad general factor of psychopathology—also known as the p factor—and two or more specific factors of psychopathology with putatively different causes and mechanisms (Caspi et al., 2014; Caspi & Moffitt, 2018; Kotov et al., 2017; Lahey et al., 2012; Lahey, Krueger, Rathouz, Waldman, & Zald, 2017; Lahey, Van Hulle, Singh, Waldman, & Rathouz, 2011).

One impediment to research on the hierarchical structure of psychopathology is the use of alternative statistical models with different interpretations to define the hierarchies. In the ‘bassackward’ strategy, a series of *separate* correlated-factors analyses are conducted (Goldberg, 2006). One factor is extracted in the first model to define the general factor. Two factors are extracted in a separate second model, followed by three factors in the next model, and so on, until all justifiable factors have been extracted. Because the only connections between levels of such hierarchies are correlations calculated after the fact between factors in the separate successive models, factors are defined in each model without parsing variance attributed to factors at other levels (Michelini et al., 2019).

Some studies use *second-order models* (Carragher et al., 2016; Sunderland et al., 2019), in which every symptom (or dimension of symptoms) loads on one of three or more correlated lower-order factors, which load on the second-order general factor (Figure 1). Thus, the general factor is defined by the variance shared by the lower-order factors. As a result, the general and lower-order factors in second-order models are not independent, but share variance. Other studies use *bifactor models* (Holzinger & Swineford, 1937; Mansolf & Reise, 2017; Reise, 2012) to define the hierarchy of psychopathology dimensions (Bloemen et al., 2018; Carragher et al., 2016; Caspi et al., 2014; Lahey et al., 2012; Lahey et al., 2015; Lahey et al., 2018; Sunderland et al., 2019). In bifactor models (Figure 1), each symptom (or dimension of symptoms) loads both on the general factor and on one of two or more orthogonal specific factors. These loadings completely and optimally partition the total variance in symptoms into separate and orthogonal general and specific factors (Mansolf & Reise, 2017; Reise, 2012).

**Figure 1.**
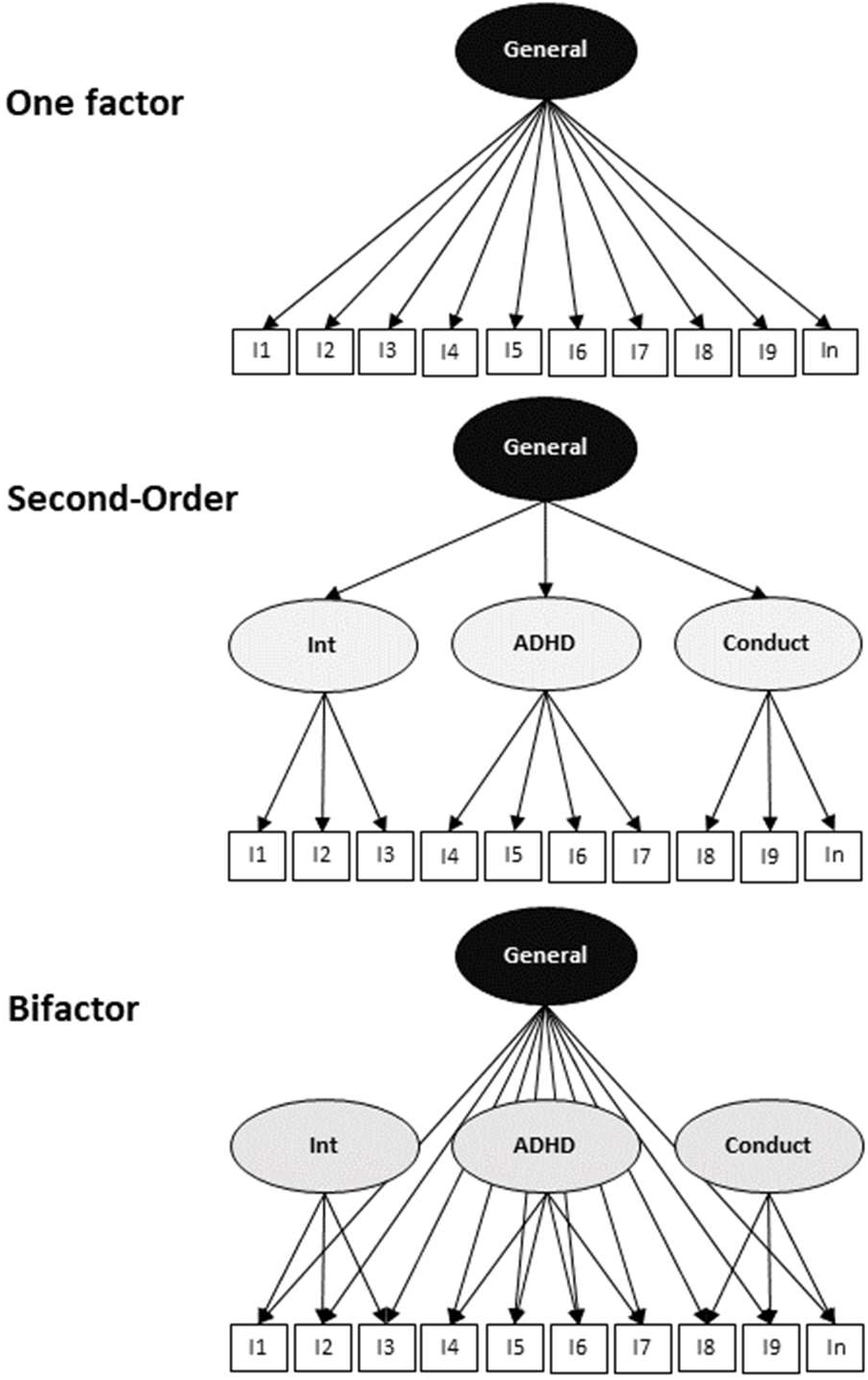
Diagrams of confirmatory factor models compared in the present analyses.

As detailed in the psychometric literature, bifactor and second-order factor models are similar on the surface, but different in interpretability (Bornovalova, Choate, Fatimah, Petersen, & Wiernik, 2020; Mansolf & Reise, 2017). Although the general factors defined in these two models are highly correlated, even the most straightforward test of criterion validity—the association between *only* the general factor and relevant external variables—has different interpretations in bifactor versus second-order models.

Furthermore, tests of criterion validity of the specific/lower-order factors in these models have different interpretations in two important ways. First, unlike bifactor models, the general factor in a second-order model shares variance with the lower-order factors. This changes the meaning of associations of the general factor with criterion variables. Second, in bifactor models, one can examine the unique associations of external variables with the specific factors while controlling for the general factor (because they are defined independently), but this cannot be done in second-order models because they are not independently defined. This is because the general factor in second-order models is defined solely by the loadings of the lower-order factors on it, making the lower- and second-order general factors (as a whole) *perfectly collinear*. One could identify the external correlates of each lower-level factor one at a time, but the meaning of those associations would be ambiguous as they could partly or wholly reflect variance that is shared with the other factors in the second-order model. Alternatively, associations of each lower-order factor with external variables could be estimated while controlling for the other lower-level factors, but the associations of all three factors with the criteria would likely be overestimated because the lower-order factors would still share variance with general factor in second-order models. If one chose instead to use the residuals (“disturbances”) of the lower-order factors (i.e., first regressing out the second-order general factor), 100% of the shared variance between the second-order general factor and each lower-order factor would be attributed to the second-order general factor, underestimating associations of lower-order factors with criteria.

In contrast, because all factors are orthogonal in bifactor models, one can simultaneously regress measured criterion variables on the general and specific factors to determine if each of these factors independently accounts for unique variance in that variable. Such advantages of bifactor models for discovering the correlates of psychopathology at each level of the hierarchy are not in dispute (Bornovalova et al., 2020; Watts, Poore, & Waldman, 2019). What is in dispute, however, is whether the psychometric properties of all factors in bifactor models are sufficient to realize its logical advantages (Conway, Mansolf, & Reise, 2019; Watts et al., 2019). This issue can be addressed in part using psychometric indices recommended for the evaluation bifactor models (Rodriguez, Reise, & Haviland, 2016): (1) H quantifies how well each latent factor is represented by the items loading on it, and is sometimes interpreted as estimating the future replicability of the factor; (2) Explained common variance (ECV) is the proportion of the total variance in all items explained by the general factor rather than the specific factors; (3) Omega estimates the proportion of total variance in the symptoms attributable to the general and specific factors together; (4) OmegaH is the proportion of total variance attributable to each general or each specific factor, by itself; (5) Factor determinacy estimates the reliability of factor scores from the correlation between a factor and the scores generated from that factor (Grice, 2001); (6) Percent uncontaminated correlations (PUC) is the percent of all correlations among symptoms attributable purely to the general factor.

Few published studies of bifactor models have reported these psychometric indices, but two papers (Conway et al., 2019; Watts et al., 2019) have raised concerns about the psychometric adequacy of specific factors in bifactor models based on some H indices below the recommended benchmark of .70 (Hancock & Mueller, 2001). Fortunately, a recent meta-analysis calculated the psychometric indices from factor loading matrices in 49 papers that had not reported them (Constantinou & Fonagy, 2019). This meta-analysis found a mean H for the general factor of 0.91 and 0.69 for specific factors. Mean factor determinacy was adequate to high for all factors, indicating that general and specific factor scores were reliable enough to test their criterion validity. Furthermore, the mean PUC of 0.67 was somewhat below the benchmark of 0.70, thought to indicate that the symptoms are essentially unidimensional, indicating that specific factors could be defined adequately. Importantly, the psychometric indices were quite heterogeneous in these 49 studies, with these indices varying with the informant on psychopathology and being stronger when more items (i.e., symptoms or dimensions) defined each factor.

In this paper, we report the results of new factor analyses of item-level data from parent ratings of child behavior and emotions as well as tests of criterion validity using both models in the Adolescent Brain Cognitive Development (ABCD) Study (Volkow et al., 2018). These head-to-head empirical comparisons are conducted following strict requirements suggested for bifactor modeling (Bonifay, Lane, & Reise, 2017; Bornovalova et al., 2020; Sellbom & Tellegen, 2019):

1. Because bifactor models tend to fit better than correlated-traits and second-order models, even when the data do not justify it (Bonifay et al., 2017), a clear theoretical basis for use of a bifactor model is required. We use bifactor models because they operationalize the causal taxonomy proposed by Lahey et al. (2017), which posits that the hierarchy of correlated phenotypes results from a hierarchy of genetic and environmental causal influences, some of which nonspecifically influence all dimensions of psychopathology to varying extents through the general factor, whereas other causal factors influence dimensions within subdomains of psychopathology (e.g., internalizing and externalizing), and still other causal factors influence only a single dimension of symptoms.
2. Samples must be at least reasonably unbiased and large enough to reliably estimate parameters and psychometric indices.
3. The general and the specific factors defined in bifactor models must be reliably measured and replicable over time in the same persons. The indices just described are useful in this regard, but psychometric indices based on a *single assessment* are not the only, and probably not the best, way to assess the reliability and stability of factors. Notably, several longitudinal studies have found that bifactor structures are replicated and each factor is correlated with itself over time in the same individuals, which could happen only if the factors were reliably measured (Castellanos-Ryan et al., 2016; Gluschkoff, Jokela, & Rosenstrom, 2019; McElroy, Belsky, Carragher, Fearon, & Patalay, 2018; Neumann et al., 2016; Olino et al., 2018; Snyder, Young, & Hankin, 2017).
4. Each factor must be valid at least in the sense of criterion validity. Notably, the general factor and specific factors of psychopathology defined in bifactor models have each already been found account for unique variance in adverse functional outcomes, including psychoactive drug prescriptions, incarceration, poor academic progress, suicidal behavior, and self-harm (Haltigan et al., 2018; Lahey et al., 2015; Pettersson, Lahey, Lundström, Larsson, & Lichtenstein, 2018; Sallis et al., 2019), and in theoretically related constructs, including trait negative emotionality (Caspi et al., 2014; Class et al., 2019) and executive functions (Bloemen et al., 2018; Martel et al., 2017; Shields, Reardon, Brandes, & Tackett, 2019). In the present analyses, we assess the criterion validity of each factor defined in bifactor and second-order models using a diverse set of theoretically and practically relevant external variables that are measured independently (i.e., without shared method variance with psychopathology).

## METHOD

### Sample

The present analyses used data from wave 1 of the ABCD Study. This sample was recruited at 22 sites across the United States at 9-10 years of age as part of a planned longitudinal study. The sites do not represent the population of the United States, but the same unbiased recruitment process was used within every site (Garavan et al., 2018) and post-stratification weights can be used to adjust the sample to be more representative (Heeringa & Berglund, 2018). Parent ratings of child psychopathology were collected (*N* = 11,866; 47.9% female). Most (8,142) participants were one child of singleton birth from different families, but 3,724 had a twin or non-twin sibling in the study. Parents classified the children as Non-Hispanic white (52.08%), Black (14.99%), Hispanic (20.29%), and other race-ethnic groups (12.63%). We conducted exploratory and preliminary analyses in one of two stratified random halves of the sample (N = 5,932 with non-missing psychopathology data), selected within data collection sites. The second random half of the sample (N = 5,934) was used to conduct planned confirmatory factor analyses that were specified using the results of the exploratory analyses.

### Measures

The Child Behavior Checklist (CBCL) (Achenbach, 2009) is a parent rating scale of child behavior consisting of 119 items describing behaviors and emotions on a scale of 0 = not true (as far as you know), 1 = somewhat or sometimes true, or 2 = very true or often true. Missing data on CBCL items was < 0.1%. We used other measures administered in the ABCD Study in tests of the criterion validity of psychopathology factors. To avoid confounding by method variance, all criterion measures were based on youth reports or formal testing of the children. Parent reports of functional impairment were the exception, but these represented decisions made by social systems (e.g., special education placement).

#### Cognitive Measures

The ABCD cognitive test battery (Luciana et al., 2018) assessed fluid reasoning, episodic memory, flexible thinking, attention, working memory, learning, mental rotation, processing speed, vocabulary comprehension, and oral reading. It included the Little Man Task (Acker & Acker, 1982), Rey Auditory Verbal Learning Task (mean total correct of trials 1-5) (Taylor, 1959), Matrix Reasoning subtest from the Wechsler Intelligence Test for Children-V (Wechsler, 2014), Picture Vocabulary Task (PPVT) (Gershon et al., 2013), Oral Reading Recognition Task (Gershon et al., 2013), Pattern Comparison Processing Speed Test (Carlozzi, Beaumont, Tulsky, & Gershon, 2015), Dimensional Change Card Sort Task (Zelazo, 2006), List Sorting Working Memory Test (Tulsky et al., 2013), Picture Sequence Memory Test (Bauer et al., 2013), and Flanker Task (Fan, McCandliss, Sommer, Raz, & Posner, 2002).

#### Dispositional Measures

The ABCD battery (Barch et al., 2018) included two measures of dispositional traits completed by child self-report: an ABCD short form of the UPPS impulsivity measure (Zapolski, Stairs, Settles, Combs, & Smith, 2010) and the prosocial subscale from the Strengths and Difficulties Questionnaire (Goodman, 1997; Hawes et al., 2019).

#### Suicide and Self-Harm Behavior

Children were administered a version of the Kiddie Schedule for Affective Disorders and Schizophrenia in which they reported if they had ever engaged in self-injurious behavior without suicidal intent, concrete suicidal ideation or planning, an interrupted, aborted, or completed suicide attempt, or planned a suicide attempt (Kaufman et al., 2016; Lisdahl et al., 2018).

#### Functional Impairment

Three measures of functional impairment included ever receiving mental health services, school detentions or suspensions, and enrollment in any form of special education, except gifted programs.

### Statistical Analyses

Missing data were minimal; only 2 of 5,936 selected for the second split sample had missing CBCL data. Among the 5,934 with non-missing CBCL data, 93.61% had non-missing data on all cognitive variables. Similarly, 99.66% had non-missing data on all dispositional scales and 96.66% had non-missing values on all impairment measures. All factor analyses and structural equation models were conducted in Mplus 8.3 using the mean- and variance-adjusted weighted least squares (WSLMV) estimator, which uses pairwise deletion for missing data (Muthén & Muthén, 1998-2017). All analyses accounted for the stratification of the sample in data collection sites, used post-stratification weights (Heeringa & Berglund, 2018), and accounted for clustering within families.

#### Exploratory Analyses in the First Half of the Sample

Exploratory analyses reduced the number of CBCL items to those most strongly associated with psychopathology at this age. The CBCL casts a broad net, including behaviors of concern to parents not typically viewed as symptoms of psychopathology, such as biting fingernails and constipation. Additionally, the CBCL includes items that are more appropriate for older ages, such as alcohol consumption and smoking. The item, “wishes to be of the opposite sex,” was eliminated because it does not reflect psychopathology. The steps in the exploratory analyses were: Eight items referring to behaviors typical of adolescents were eliminated because ratings above 0 were < 0.5%, or were < 1.0% *and* it was not possible to estimate polychoric correlations with other items (Table S1). Three pairs of items that referenced similar behaviors correlated >.85 were combined in composites by taking the mean rating of the items and rounding to achieve 0, 1, 2 scoring (Table S1). Exploratory structural equation (ESEM) models (Asparouhov & Muthén, 2009) were conducted with OBLIMIN rotation. Parallel analysis (Horn, 1965) with Glorfeld correction (Glorfeld, 1995) indicated that up to 9 factors could be extracted (Figure S1). The minimum average partial criterion (Velicer, 1976) indicated that up to 6 factors could be extracted (Figure S1). We extracted four interpretable factors and all CBCL items with a loading ≥ 0.40 on at least one factor were retained. Retained items were subjected to ESEM specifying 2, 3, or 4 correlated factors in the first half of the sample (Tables S2-S4).

#### Confirmatory Analyses in the Second Half of the Sample

Bifactor and second-order confirmatory models were specified based on the results of the ESEMs. Items loading >0.40 on two factors were assigned to the factor with the higher loading. As required for bifactor models, each item loaded on the general factor and only one specific factor. All other loadings were fixed to zero, and all factors were specified to be orthogonal (Reise, 2012).

#### Comparisons of the Models

We directly compared the bifactor and second-order models specifying a general factor and three specific or three lower-order factors of psychopathology in two ways. First, estimated factor scores from Mplus were plotted. Second, tests of criterion validity were conducted using structural equation modeling in Mplus. For the bifactor model, each measured criterion variable was regressed simultaneously in these SEMs on the general and specific psychopathology factors. Probit regressions were estimated in SEM for binary criterion variables. Based on previous studies that identified a general factor of cognitive ability test scores using similar test batteries in children (Friedman & Miyake, 2017; Martel et al., 2017), the substantially correlated cognitive measures defined a single latent factor in a confirmatory measurement model.

The strategy for testing the criterion validity of the second-order model was necessarily different. Because the second-order general factor is defined by the lower-order factors, it was necessary to conduct two separate regression analyses (within the SEM), one for only the general factor and one in which criteria were simultaneously regressed on lower-order internalizing, conduct problems, and ADHD factors. For both bifactor and second-order models, covariates of no interest (child’s age in months, sex, and race-ethnicity) were included in the models. *Sensitivity Analysis*

To evaluate the impact of missing data on cognitive test measures, we first used listwise deletion to drop all participants with missing data on any measure. Second, we imputed missing data once for cognitive measures and using SAS PROC MI.

## Results

### ESEMs Based in the First Half of the Sample

Factor loadings for 2-, 3-, and 4-factor ESEMs of the retained CBCL items are in Tables S2-S4. When three factors were extracted, the items defined factors of internalizing (i.e., fears, depression, insecurity, and somatic complaints), conduct problems (i.e., oppositional and conduct disorder), and attention-deficit/hyperactivity disorder (ADHD) and neurodevelopmental problems (e.g., immaturity, poor coordination, and sluggish cognitive tempo), all of which have long been known to be correlated (Achenbach et al., 1989; Quay, 1986). Items referring to strange ideas and behaviors loaded on the ADHD factor. When a fourth factor was extracted, somatic complaint items split from the internalizing factor.

### Confirmatory Factor Analyses in the Second Half of the Sample

Results for a bifactor model with two specific factors are in Table S5. We focused on bifactor and second-order models with three or four specific/lower-order factors, however, which allow second-order models to be identified. Factor loadings from models with three specific/lower-order factors are in Tables S6 and S7 (Tables S8 and S9 for models specifying four specific/lower-order factors). Each model fit the data well (Table S10). Because of difficulties in choosing between substantively different models that fit the data well using fit statistics, we evaluated the two structural models in terms of their criterion validity (Bonifay et al., 2017; Greene et al., 2019).

As shown in Table 1, all factors in both models had acceptable H indices >0.70 (Hancock & Mueller, 2001) and each specific factor in the bifactor models was reliable according to omega statistics. ECV and omegaH indicated that the general factor is robust and explains the majority of the estimated reliability of each specific factor. Nonetheless, the factor determinacy indices show that each of the specific factors is sufficiently determined to calculate specific factor scores that are orthogonal to each other and to the general factor score. This means that any unique associations of a specific factor score with external variables can be interpreted as unique to that factor and can be used to evaluate their criterion validity and examine associations with variables relevant to causes and mechanisms.

**Table 1.**
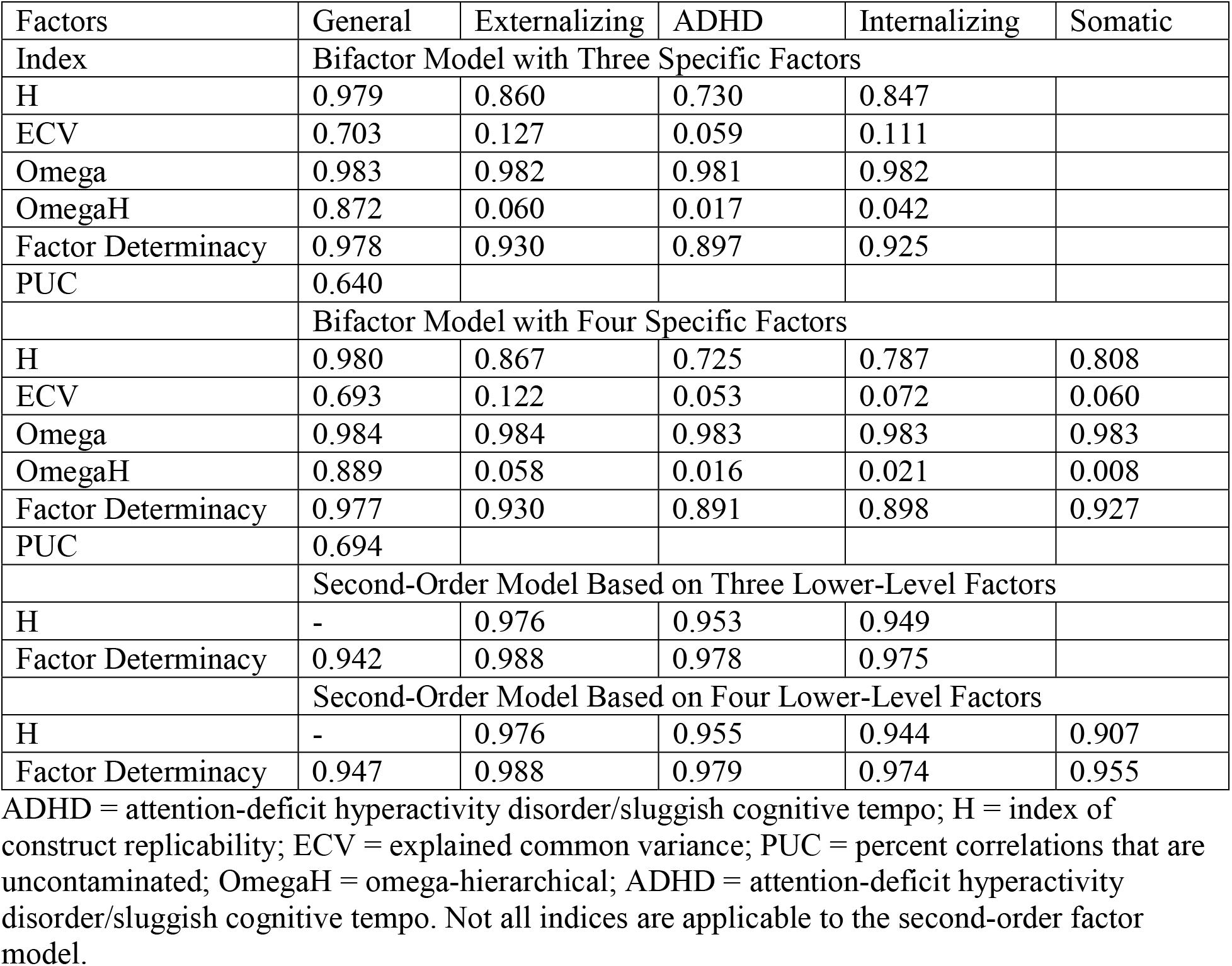
Psychometric indices for each latent factor defined in bifactor and second-order model specifying three or four specific/lower-level in the second split half of the ABCD Study sample (N = 5926) to which the statistic is applicable.

### Correlations between Factors across Models

As shown in Figure 2, estimated factor scores for the general factors defined in the two models were highly correlated. In contrast, the specific internalizing, conduct, and ADHD factor scores were moderately correlated across models. The relationships in Figure 2 are more complex than they first appear, however. Because general and specific/lower-order factor scores are independent in bifactor models, as shown in Figures S3 and S4, individuals high on ADHD, for example, are often low on the general factor and vice versa. In contrast, that is not the case in second-order models because these factors are highly correlated. As a result, Figure 2 reveals an apparent Simpson’s Paradox (Simpson, 1951) in which the overall *positive linear* correlations between specific and lower-order scores across models obscure the fact that, at low values, ADHD scores, for example, from the two models are *negatively correlated*. This occurs because at higher general factor scores, ADHD scores in both models tend to be high, but when general factor scores are low, ADHD scores are lower in second-order than bifactor models. This negative correlation in the lower range is one of the ways the bifactor model achieves specific factor scores orthogonal to the general factor when considered across the full range of scores.

**Figure 2.**
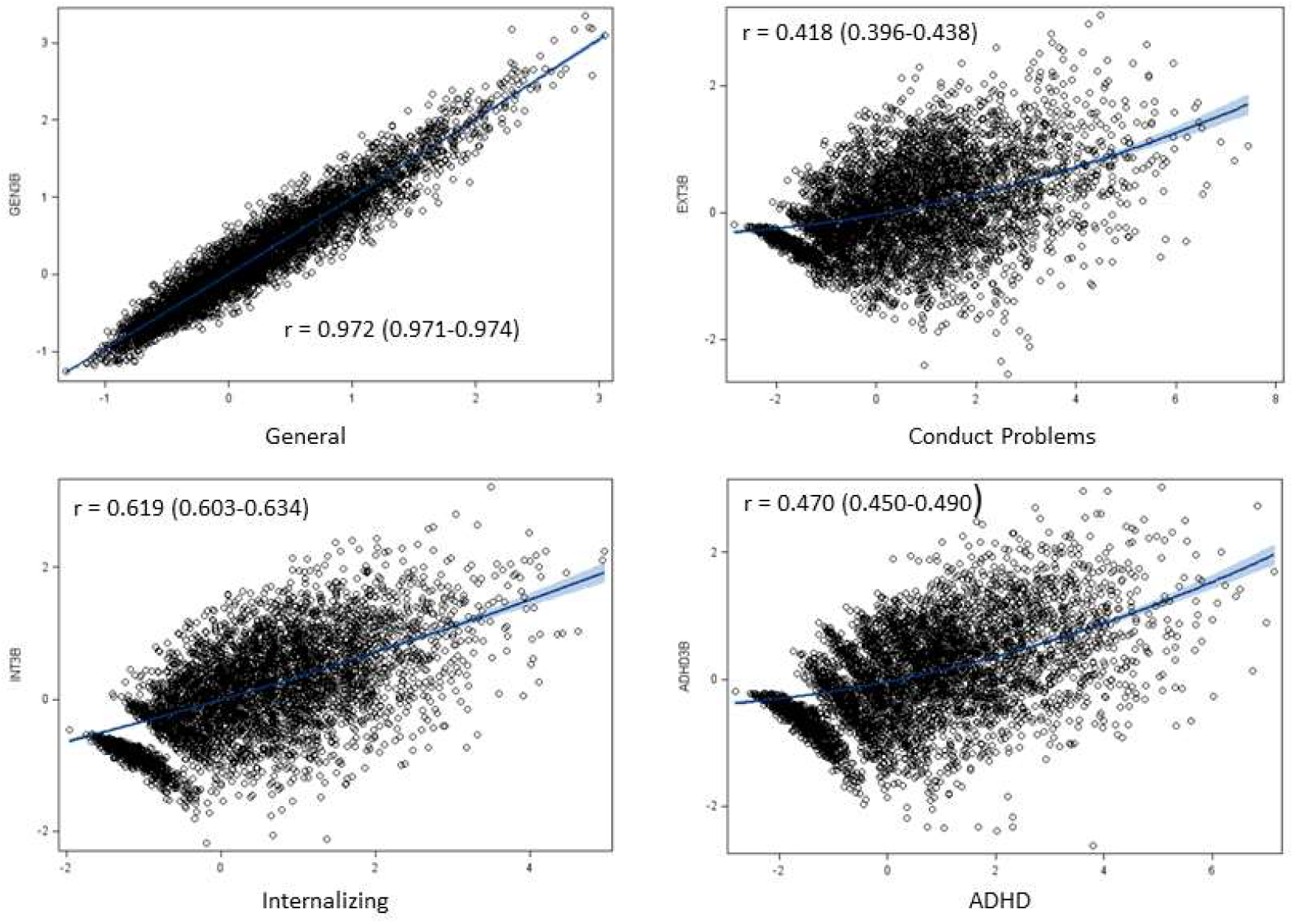
Scatterplots and correlations between estimated general and specific or lower-order factor scores defined in bifactor (vertical axes) and second-order confirmatory factor models (horizontal). Blue shading defines the 95% confidence interval of the best-fit regression line.

### Criterion Validity

Simultaneous regressions within the SEMs revealed that each of the latent general and specific factors defined by the bifactor model explained significant unique variance after correction for multiple testing in the criterion variables (Table 2). The latent second-order general factor was significantly associated with the same criterion variables as in the bifactor models (Table 3) and each of the three lower-level factors explained significant unique variance in multiple criterion variables, controlling for the variance explained by the other two lower-order factors. The results for the bifactor model with four specific factors and the second-order model based on four lower-order factors presented in Tables S11 and S12 were virtually identical, except for the addition of the somatic complaints factor. Sensitivity analyses based on listwise deletion and imputation of missing cognitive test scores yielded similar results (Table S13).

**Table 2.** Results of simultaneous regressions (within an SEM) of each independently measured criterion variable on the latent general factor and three (or four) specific factors defined in bifactor models and on demographic covariates of no interest (age, sex, and race-ethnicity) in the second split half of the ABCD Study sample (N = 5926).

| Bifactor Model with Three Specific Factors |  |  |  |  |  |  |  |  |
| --- | --- | --- | --- | --- | --- | --- | --- | --- |
| Criterion Variable | Factors |  |  |  |  |  |  |  |
|  | General |  | Specific Internalizing |  | Specific Conduct |  | Specific ADHD |  |
| | $\beta$ | P < | $\beta$ | P < | B | P < | B | P < |
| *Functional Impairment (Binary)* |  |  |  |  |  |  |  |  |
| Detention/suspension | **0.380** | **0.001** | **-0.084** | **0.011** | **0.409** | **0.001** | 0.040 | 0.307 |
| Mental health service | **0.577** | **0.001** | **0.131** | **0.001** | **0.106** | **0.001** | 0.064 | 0.040 |
| Special Education (Behavior) | **0.295** | **0.001** | 0.024 | 0.704 | **0.253** | **0.001** | 0.093 | 0.112 |
| Special Education (Learning) | **0.288** | **0.001** | 0.003 | 0.956 | 0.059 | 0.212 | **0.187** | **0.004** |
| *Youth-Reported Harmful Behaviors (Binary)* |  |  |  |  |  |  |  |  |
| Suicidal Behavior | **0.270** | **0.001** | 0.032 | 0.456 | 0.098 | 0.072 | -0.002 | 0.979 |
| Non-suicidal Self Harm | **0.220** | **0.001** | 0.057 | 0.098 | 0.081 | 0.076 | **0.093** | **0.025** |
| *Youth-Rated Dispositional Constructs (Standardized Continuous)* |  |  |  |  |  |  |  |  |
| UPPS low premeditation | **0.132** | **0.001** | **-0.079** | **0.001** | **0.099** | **0.001** | **0.109** | **0.001** |
| UPPS low perseverance | **0.128** | **0.001** | 0.018 | 0.310 | **0.075** | **0.001** | **0.248** | **0.001** |
| UPPS negative urgency | **0.172** | **0.001** | **-0.059** | **0.002** | **0.087** | **0.001** | -0.001 | 0.966 |
| UPPs positive urgency | **0.113** | **0.001** | **-0.089** | **0.001** | **0.097** | **0.001** | **0.085** | **0.001** |
| UPPS sensation seeking | -0.005 | 0.796 | **-0.051** | **0.007** | **0.104** | **0.001** | 0.034 | 0.169 |
| Prosociality | **-0.073** | **0.001** | 0.033 | 0.088 | **-0.065** | **0.008** | -0.030 | 0.211 |
| *Latent Cognitive Test Scores* |  |  |  |  |  |  |  |  |
| Executive functioning | **-0.091** | **0.001** | 0.021 | 0.293 | **-0.241** | **0.001** | **-0.328** | **0.001** |

Coefficients in bold are significant after FDR correction (adopting a 5% false discovery rate) for 52 tests. ADHD = attention-deficit hyperactivity disorder/sluggish cognitive tempo. Continuous dispositional and executive functioning measures were normalized to a mean of 0 and standard deviation of 1.

**Table 3.** Results of regressions of each independently measured criterion variable on only the latent general factor and demographic covariates of no interest (age, sex, and race-ethnicity) and in separate models simultaneously on only the three lower-order factors defined in second-order factor models and on demographic covariates of no interest (age, sex, and race-ethnicity) in the second split half of the ABCD Study sample (N = 5926).

| Second-Order Model Based on Three Lower-Order Factors |  |  |  |  |  |  |  |  |
| --- | --- | --- | --- | --- | --- | --- | --- | --- |
| Criterion Variable | Factors |  |  |  |  |  |  |  |
|  | General |  | Lower-Order Internalizing |  | Lower-Order Conduct |  | Lower-Order ADHD |  |
| | B | P < | $\beta$ | P < | $\beta$ | P < | $\beta$ | P < |
| Regressions of Criterion Variables on Only the Second-Order General Factor and Covariates |  |  |  |  |  |  |  |  |
| *Functional Impairment (Binary)* |  |  |  |  |  |  |  |  |
| Detention/suspension | **0.495** | **0.001** |  |  |  |  |  |  |
| Mental health service | **0.634** | **0.001** |  |  |  |  |  |  |
| Special educ (Behavior) | **0.381** | **0.001** |  |  |  |  |  |  |
| Special educ (Learning) | **0.332** | **0.001** |  |  |  |  |  |  |
| *Youth-Reported Harmful Behaviors (Binary)* |  |  |  |  |  |  |  |  |
| Suicidal behavior | **0.301** | **0.001** |  |  |  |  |  |  |
| Non-suicidal self harm | **0.263** | **0.001** |  |  |  |  |  |  |
| *Youth-Rated Dispositional Constructs (Standardized Continuous)* |  |  |  |  |  |  |  |  |
| UPPS low premeditation | **0.161** | **0.001** |  |  |  |  |  |  |
| UPPS low perseverance | **0.188** | **0.001** |  |  |  |  |  |  |
| UPPS negative urgency | **0.184** | **0.001** |  |  |  |  |  |  |
| UPPS positive urgency | **0.135** | **0.001** |  |  |  |  |  |  |
| UPPS sensation seeking | 0.017 | 0.344 |  |  |  |  |  |  |
| Prosociality | **-0.088** | **0.001** |  |  |  |  |  |  |
| *Latent General Executive Functioning Test Scores* |  |  |  |  |  |  |  |  |
| Cognitive functioning | **-0.197** | **0.001** |  |  |  |  |  |  |
| Simultaneous Regressions of Criterion Variables on Only the Three Lower-Order Factors and Covariates |  |  |  |  |  |  |  |  |
| *Functional Impairment (Binary)* |  |  |  |  |  |  |  |  |
| Detention/suspension |  |  | **-0.218** | **0.001** | **0.683** | **0.001** | -0.020 | 0.705 |
| Mental health service |  |  | **0.247** | **0.001** | **0.251** | **0.001** | **0.170** | **0.001** |
| Special Educ (Behavior) |  |  | -0.027 | 0.762 | **0.372** | **0.001** | 0.027 | 0.748 |
| Special Educ (Learning) |  |  | -0.017 | 0.843 | 0.028 | 0.758 | **0.332** | **0.001** |
| *Youth-Reported Harmful Behaviors (Binary)* |  |  |  |  |  |  |  |  |
| Suicidal Behavior |  |  | 0.083 | 0.245 | **0.174** | **0.032** | 0.053 | 0.498 |
| Non-suicidal Self Harm |  |  | 0.089 | 0.101 | 0.063 | 0.351 | 0.125 | 0.044 |
| *Youth-Rated Dispositional Constructs (Standardized Continuous)* |  |  |  |  |  |  |  |  |
| UPPS low premeditation |  |  | **-0.143** | **0.001** | **0.091** | **0.001** | **0.203** | **0.001** |
| UPPS low perseverance |  |  | -0.052 | 0.059 | **-0.098** | **0.002** | **0.346** | **0.001** |
| UPPS negative urgency |  |  | **-0.070** | **0.011** | **0.211** | **0.001** | 0.034 | 0.312 |
| UPPs positive urgency |  |  | **-0.168** | **0.001** | **0.137** | **0.001** | **0.151** | **0.001** |
| UPPS sensation seeking |  |  | **-0.125** | **0.001** | **0.109** | **0.002** | 0.018 | 0.614 |
| Positive emotionality |  |  | **-0.093** | **0.007** | 0.048 | 0.092 | -0.038 | 0.269 |
| *Latent General Executive Functioning Test Scores* |  |  |  |  |  |  |  |  |
| Cognitive functioning |  |  | **0.209** | **0.001** | 0.065 | 0.052 | **-0.327** | **0.001** |

Coefficients in bold are significant after FDR correction (adopting a 5% false discovery rate) for 52 tests. ADHD = attention-deficit hyperactivity disorder/sluggish cognitive tempo. Continuous dispositional and executive functioning measures were normalized to a mean of 0 and standard deviation of 1.

## DISCUSSION

This study provides new information on two alternative statistical models of the hierarchical dimensional structure of psychopathology using data from the largest study of child psychopathology to date. Both bifactor and second-order factor models of CBCL symptoms fit the data well. The psychometric indices reported in Table 1 provide strong support for the general factor in both models; indeed, they raise the radical possibility that psychopathology measured by the CBCL at this age is unidimensional (i.e., adequately captured by the general factor alone). The omegaH (~0.87) and ECV (~0.70) indicated that the general factor is strong, particularly when considered with the PUC, which indicates that 64% of item correlations are explained by the general factor. Furthermore, when omega and omegaH are compared, >90% of the reliability of the specific factor scores (e.g., specific conduct problems) is due to the general factor. It should be noted, however, that a study of psychopathology in older youth (Sunderland et al., 2019) found far less evidence of uni-dimensionality.

Moreover, other results of the present analyses argue against uni-dimensionality and support the *multi-dimensionality* of parent-rated psychopathology. Like Sunderland et al. (2019), the present bifactor and second-order models both defined all factors with strong determinacy and acceptable construct replicability (Table 1). Therefore, it was possible to conduct critical tests of the criterion validity to determine if the lower-level/specific factors are valid.

### Criterion Validity of General and Specific/Lower-Order Factors

Like previous studies (Lahey et al., 2017; Pettersson et al., 2018), the general factor defined in the bifactor model accounted for substantial unique variance in external criteria. For example, as presented in Table 2, each 1 SD increase in general factor scores was associated with a .38 SD (+- 0.03) greater likelihood of being detained or suspended from school for misbehavior, over and above demographic covariates and the robust association with specific conduct problems and the inverse association with internalizing psychopathology. Furthermore, *only* the general factor was significantly correlated with youth-reported suicidal planning and attempts, and each 1 SD increase in general factor scores was associated with a .27 SD (+- 0.04) greater likelihood of engaging in non-suicidal self-injury. The general factor in the bifactor model also was significantly associated with obtaining mental health and special education services. These associations suggest the potential clinical utility of the general factor in evaluating prognostic risk in children.

At a theoretical level, it is important that the results in Table 2 replicate previous associations of the bifactor general factor with independent measures of cognitive functioning (Martel et al., 2017; Shields et al., 2019) and negative emotionality (Caspi & Moffitt, 2018; Lahey et al., 2017). Although negative emotionality was not measured in this study, UPPS negative urgency was measured, which is correlated with negative emotionality (Settles et al., 2012). Thus, the present findings that the general factor was significantly associated with both cognitive functioning and UPPS negative urgency are consistent with the hypothesis that lower executive functioning and higher negative emotionality are two processes that nonspecifically contribute to risk for all forms of psychopathology, which is captured by the general factor (Lahey et al., 2017). Additionally, the finding that both positive and negative urgency are associated with the general factor in the same direction is consistent with the hypothesis that the general factor reflects impulsive (i.e., unregulated) responsivity to both positive and negative emotions (Carver, Johnson, & Timpano, 2017).

Associations of the specific factors defined in the bifactor model with criterion variables reported in Table 2 replicate previous studies (reviewed by Lahey et al., 2017). Consistent with the meanings of these factors, the specific internalizing and conduct problems factors were each positively associated with mental health service, but associated with school detentions and expulsions, and with premeditation, negative and positive urgency, and sensation seeking in opposite directions as would be expected. The specific ADHD factor exhibited criterion validity in its associations with risk for special education placement for learning problems and non-suicidal self-harm, with three dispositional constructs theoretically associated with ADHD (low premeditation, low perseverance, and high positive urgency), and in its robust inverse association with executive functioning. Considered together, these robust associations with independently measured criterion variables support the validity of both the general and the three specific factors defined in the bifactor model.

The results in Table 3 similarly support the criterion validity of psychopathology factors defined in the second-order model. The *pattern* of significant and nonsignificant associations is similar, but the *interpretations* of these associations are different. Because the second-order general factor is defined by the loadings of lower-order factors on it, it was not possible to simultaneously regress criterion variables on both the general and the lower-order factors to test each factor’s unique criterion validity. Therefore, tests of criterion validity of lower-order factors and the second-order general factor had to be conducted separately—identifying the unique correlates of the lower-level factors while controlling for the second-order general factor is impossible. It would be analogous to including three items in a predictive model along with the sum of those same three items.

Notably, the latent general factor defined in the second-order model explained an average of 4.9% more variance in these criterion variables (range 1.2 – 11.5%; median 4.3%) than the general factor in the bifactor model. This does not mean that the second-order model is superior to the bifactor model, however. Indeed, the apparent superiority of the second-order model *reflects its major limitation* for research on variables related to the causes and mechanisms of each factor of psychopathology. The stronger associations of the second-order general factor with criterion variables are due to *contamination* of the general factor and lower-order factors with each other in the second-order model. For example, the association of the second-order general factor with school detentions/suspensions is inflated because the general factor contains variance associated with the lower-order conduct problems factor, which is strongly associated with detentions/suspensions. We can see this at the top of Table 2 where the total variance explained in detentions/suspensions by the general and specific factors in the bifactor model is 0.380^2^ + (−0.084)^2^ + 0.409^2^ + 0.040^2^ = 0.32, with half of that variance coming from conduct problems (0.409). The variance associated with conduct problems is not included in the estimated association of the general factor with detentions/suspensions in the bifactor model, but it spuriously boosts the variance explained by the general factor in the second-order model. This difference in interpretation of such associations is a key difference between the models.

### Choosing between Bifactor and Higher-Order Models of Psychopathology

Given that all latent factors defined in both bifactor and second-order models were well-defined and valid, the choice between these two models depends on its intended use. The bifactor model is useful for testing models that posit that some risk factors and psychobiological mechanisms are common to all forms of psychopathology, whereas other risk factors are only related to specific factors (Caspi et al., 2014; Caspi & Moffitt, 2018; Lahey et al., 2017; Lahey et al., 2011). This is because, in a bifactor model, the variance in symptoms is empirically partitioned among the general and specific factors (Mansolf & Reise, 2017; Reise, 2012). Thus, the bifactor model is useful when the goal is to identify both common and *unique* correlates of the dimensions of psychopathology.

### Limitations and Implications for Future Research

It should be emphasized that the results of any factor analysis reflect the items that are analyzed and the sample in which they are analyzed. We restate this obvious fact to guard against the reification of constructs such as the general factor of psychopathology. This admonition is not unique to bifactor models, but applies to any factor analysis. There are several important implications of the present findings for future studies, especially in the context of previous research. These findings indicate that the factors defined in the bifactor model based on CBCL items have sufficient psychometric properties (e.g., determinacy) to allow the advantages of the orthogonal factors in bifactor models to be realized in discovery studies, including the important longitudinal ABCD Study. This means, for example, that the bifactor model can be used to test associations of orthogonal factors of psychopathology to polygenic risk scores in this sample. The general factor has already been shown to be moderately heritable and significantly related to such independently measured genetic risk scores (Allegrini et al., 2020; Neumann et al., 2016), but we need to learn much more. Similarly, well-powered analyses of ABCD structural and functional neuroimaging data will not only help reveal important psychobiological mechanisms related to psychopathology risk in specific and transdiagnostic ways, but could further support the validity of factor scores defined in bifactor models (Bornovalova et al., 2020).

Note that the present findings support such research in the ABCD Study, but do not mean that bifactor models of psychopathology will always have adequate psychometric properties; as with any factor model, that needs to be demonstrated on a study by study basis. Adequate psychometric properties flow from adequate methodology. In a broader sense, the most important obstacle to the field is that no study of the hierarchy of psychopathology has used an instrument that comprehensively measures all symptoms of all forms of psychopathology, including what were previously referred to as clinical and personality disorders (Forbes et al., 2017). If such a comprehensive measure is successfully developed and validated, we will finally be in a position to define the hierarchical structure of psychopathology and to discover its correlates and causes in future studies. One very helpful but inexpensive step towards this goal would be to supplement the CBCL with additional items in future waves of the ABCD Study.

It is important to note the implications of one current finding for the measurement of psychopathology in future waves of the ABCD Study using the CBCL. It was not possible to include a number of CBCL items in the present analyses due to their low endorsements at ages 9-10 years. These items refer to symptoms that are centrally important to the ABCD Study such as psychotic symptoms, drinking, and substance use, but are uncommon at this age. As the ABCD sample ages, however, they will become more prevalent. This means that it likely will not be possible to base factor analyses on the same subset of items at different ages. A valid way to characterize both stability and developmental change in psychopathology in the ABCD sample must be developed.

