## Supplemental Tables and Figure for "Criterion Validity and Relationships between Alternative Hierarchical Dimensional Models of General and Specific Psychopathology"

Table S1. Child Behavior Checklist (CBCL) items eliminated or combined in composites before final confirmatory factor analyses using data from the first split half of the wave 1 ABCD Study sample (N = 5932 with nonmissing CBCL data).

Step 1. Eight items were eliminated if their endorsements above 0 were < 0.5%, or if their endorsements above 0 were < 1.0% and was not possible to estimate a polychoric correlation between that item and at least one other item:

| | |
|---|---|
| cbcl_q02 | alcohol |
| cbcl_q59 | plays with own sex parts in public |
| cbcl_q67 | runs away from home |
| cbcl_q73 | sexual problems |
| cbcl_q96 | thinks about sex too much |
| cbcl_q99 | tobacco |
| cbcl_q101 | truancy |
| cbcl_q105 | drugs |

Step 2. Three pairs of items that reference similar behaviors and whose polychoric correlations were >.85 were combined in composites by taking the mean rating of the correlated items and rounding the mean to achieve 0, 1, 2 scoring:

| Composite item | Original items combined | |
|---|---|---|
| Destroys things | cbcl_q20 | Destroys his/her own things |
| | cbcl_q21 | Destroys things belonging to his/her family or others |
| Inattentive | cbcl_q08 | Can't concentrate, can't pay attention for long |
| | cbcl_q78 | Inattentive or easily distracted |
| Overweight | cbcl_q53 | Overeating |
| | cbcl_q5 | Overweight |

| Table S2. Second exploratory 2-factor analysis in ESEM of the reduced set of CBCL items after the first SEM I Table S2 in the first half of the sample. | | | |
|---|---|---|---|
| Item | Brief wording | Ext | Int |
| **1** | **Acts too young for age** | **0.464** | 0.178 |
| **3** | **Argues a lot** | **0.710** | 0.090 |
| **4** | **Fails to finish things** | **0.564** | 0.211 |
| **5** | **There is very little enjoys** | 0.351 | **0.401** |
| **7** | **Bragging, boasting** | **0.528** | 0.072 |
| **9** | **Can't get mind off certain thoughts; obsessions** | **0.402** | **0.400** |
| **10** | **Can't sit still, restless, or hyperactive** | **0.713** | 0.039 |
| 13 | Confused or seems to be in a fog | 0.356 | 0.383 |
| **15** | **Cruel to animals** | **0.535** | 0.096 |
| **16** | **Cruelty, bullying, or meanness to others** | **0.769** | -0.006 |
| 17 | Daydreams or gets lost in thoughts | 0.314 | 0.356 |
| **19** | **Demands a lot of attention** | **0.584** | 0.225 |
| **22** | **Disobedient at home** | **0.843** | -0.054 |
| **23** | **Disobedient at school** | **0.894** | -0.216 |
| **25** | **Doesn't get along with other kids** | **0.637** | 0.151 |
| **26** | **Doesn't seem to feel guilty after misbehaving** | **0.772** | -0.081 |
| **27** | **Easily jealous** | **0.499** | 0.246 |
| **28** | **Breaks rules at home, school or elsewhere** | **0.927** | -0.145 |
| **30** | **Fears going to school** | 0.097 | **0.559** |
| **31** | **Fears might think or do something bad** | 0.085 | **0.583** |
| **32** | **Feels has to be perfect** | -0.129 | **0.649** |
| **33** | **Feels or complains that no one loves them** | 0.336 | **0.484** |
| **34** | **Feels others are out to get** | **0.411** | **0.448** |
| **35** | **Feels worthless or inferior** | 0.190 | **0.670** |
| 36 | Gets hurt a lot, accident prone | 0.259 | 0.309 |
| **37** | **Gets in many fights** | **0.757** | -0.069 |
| **39** | **Hangs around with others who get in trouble** | **0.614** | -0.059 |
| **41** | **Impulsive or acts without thinking** | **0.749** | 0.099 |
| **42** | **Would rather be alone than with others** | 0.139 | **0.463** |
| **43** | **Lying or cheating** | **0.745** | -0.042 |
| 46 | Nervous movements or twitching | 0.360 | 0.328 |
| **50** | **Too fearful or anxious** | 0.042 | **0.708** |
| **51** | **Feels dizzy or lightheaded** | -0.030 | **0.598** |
| **52** | **Feels too guilty** | -0.018 | **0.732** |
| **56A** | **Aches or pains** | -0.027 | **0.475** |
| **56B** | **Headaches** | -0.047 | **0.524** |
| **56C** | **Nausea, feels sick** | -0.217 | **0.810** |
| **56F** | **Stomachaches** | -0.167 | **0.709** |
| **56G** | **Vomiting, throwing up** | -0.192 | **0.581** |
| **56H** | **Other physical problems** | 0.020 | **0.444** |

| | | | |
|---|---|---|---|
| **57** | **Physically attacks people** | **0.741** | 0.005 |
| **61** | **Poor school work** | **0.615** | 0.073 |
| 62 | Poorly coordinated or clumsy | 0.337 | 0.372 |
| **65** | **Refuses to talk** | 0.263 | **0.441** |
| **66** | **Repeats certain acts over and over; compulsions** | **0.449** | 0.346 |
| **68** | **Screams a lot** | **0.635** | 0.162 |
| **69** | **Secretive, keeps things to self** | 0.291 | **0.405** |
| **71** | **Self-conscious or easily embarrassed** | 0.056 | **0.668** |
| **72** | **Sets fires** | **0.647** | -0.171 |
| **74** | **Showing off or clowning** | **0.647** | -0.020 |
| **75** | **Too shy or timid** | -0.102 | **0.595** |
| **80** | **Stares blankly** | **0.400** | 0.368 |
| **81** | **Steals at home** | **0.783** | -0.103 |
| **82** | **Steals outside the home** | **0.796** | -0.125 |
| **84** | **Strange behavior** | **0.514** | 0.309 |
| **85** | **Strange ideas** | **0.419** | 0.335 |
| **86** | **Stubborn, sullen, or irritable** | **0.567** | 0.288 |
| **87** | **Sudden changes in mood or feelings** | **0.510** | 0.386 |
| **88** | **Sulks a lot** | 0.393 | **0.460** |
| **89** | **Suspicious** | **0.493** | 0.329 |
| **90** | **Swearing or obscene language** | **0.649** | 0.026 |
| **93** | **Talks too much** | **0.461** | 0.139 |
| **94** | **Teases a lot** | **0.649** | 0.051 |
| **95** | **Temper tantrums or hot temper** | **0.676** | 0.166 |
| **97** | **Threatens people** | **0.734** | 0.111 |
| **102** | **Underactive, slow moving, or lacks energy** | 0.167 | **0.519** |
| **103** | **Unhappy, sad, or depressed** | 0.243 | **0.610** |
| **106** | **Vandalism** | **0.868** | -0.109 |
| 109 | Whining | 0.399 | 0.322 |
| **111** | **Withdrawn, doesn't get involved with others** | 0.171 | **0.625** |
| **112** | **Worries** | -0.020 | **0.758** |
| **8,78** | **Inattentive*** | **0.698** | 0.135 |
| **20,21** | **Destroys things*** | **0.739** | 0.048 |

Latent factor correlations:
Internalizing with externalizing: 0.491

Items in bold retained for the next round of exploratory analysis using data from the first random split half of the sample to specify which items loading on the specified factors in confirmatory factor analyses in the second random split half of the sample.

Table S3. Second exploratory 3-factor analysis in ESEM of the reduced set of CBCL items after the first SEM I Table S2 in the first half of the sample.

| Item | Brief wording | Ext | ADHD | Int |
|---|---|---|---|---|
| **1** | **Acts too young for age** | 0.184 | **0.463** | 0.057 |
| **3** | **Argues a lot** | **0.764** | -0.033 | 0.108 |
| **4** | **Fails to finish things** | 0.198 | **0.586** | 0.063 |
| 5 | There is very little enjoys | 0.277 | 0.221 | 0.332 |
| **7** | **Bragging, boasting** | **0.493** | 0.097 | 0.050 |
| **9** | **Can't get mind off certain thoughts; obsessions** | 0.122 | **0.505** | 0.264 |
| **10** | **Can't sit still, restless, or hyperactive** | 0.244 | **0.687** | -0.131 |
| **13** | **Confused or seems to be in a fog** | -0.075 | **0.710** | 0.185 |
| **15** | **Cruel to animals** | **0.547** | 0.022 | 0.099 |
| **16** | **Cruelty, bullying, or meanness to others** | **0.853** | -0.118 | 0.048 |
| **17** | **Daydreams or gets lost in thoughts** | -0.152 | **0.725** | 0.170 |
| **19** | **Demands a lot of attention** | **0.472** | 0.248 | 0.159 |
| **22** | **Disobedient at home** | **0.859** | -0.006 | -0.036 |
| **23** | **Disobedient at school** | **0.701** | 0.278 | -0.269 |
| **25** | **Doesn't get along with other kids** | **0.581** | 0.145 | 0.119 |
| **26** | **Doesn't seem to feel guilty after misbehaving** | **0.694** | 0.128 | -0.097 |
| **27** | **Easily jealous** | **0.526** | 0.041 | 0.231 |
| **28** | **Breaks rules at home, school or elsewhere** | **0.848** | 0.119 | -0.152 |
| **30** | **Fears going to school** | 0.121 | 0.105 | **0.509** |
| **31** | **Fears might think or do something bad** | 0.054 | 0.189 | **0.510** |
| **32** | **Feels has to be perfect** | -0.044 | 0.026 | **0.612** |
| **33** | **Feels or complains that no one loves them** | **0.408** | 0.023 | **0.461** |
| **34** | **Feels others are out to get** | **0.401** | 0.138 | 0.399 |
| **35** | **Feels worthless or inferior** | 0.166 | 0.205 | **0.589** |
| 36 | Gets hurt a lot, accident prone | 0.042 | 0.393 | 0.199 |
| **37** | **Gets in many fights** | **0.785** | -0.040 | -0.035 |
| **39** | **Hangs around with others who get in trouble** | **0.508** | 0.172 | -0.092 |
| **41** | **Impulsive or acts without thinking** | **0.450** | 0.493 | -0.025 |
| 42 | Would rather be alone than with others | 0.049 | 0.245 | 0.385 |
| **43** | **Lying or cheating** | **0.644** | 0.175 | -0.075 |
| **46** | **Nervous movements or twitching** | 0.074 | **0.495** | 0.194 |
| **50** | **Too fearful or anxious** | -0.056 | 0.307 | **0.604** |
| **51** | **Feels dizzy or lightheaded** | 0.001 | 0.102 | **0.541** |
| **52** | **Feels too guilty** | -0.019 | 0.176 | **0.655** |
| **56A** | **Aches or pains** | 0.027 | 0.041 | **0.437** |
| **56B** | **Headaches** | 0.090 | -0.062 | **0.505** |
| **56C** | **Nausea, feels sick** | 0.041 | -0.157 | **0.783** |
| **56F** | **Stomachaches** | 0.048 | -0.125 | **0.684** |
| **56G** | **Vomiting, throwing up** | -0.005 | -0.116 | **0.559** |
| 56H | Other physical problems | 0.027 | 0.100 | 0.395 |
| **57** | **Physically attacks people** | **0.827** | -0.126 | 0.065 |
| **61** | **Poor school work** | 0.245 | **0.571** | -0.072 |
| **62** | **Poorly coordinated or clumsy** | -0.028 | **0.612** | 0.204 |

| | | | | |
|---|---|---|---|---|
| 65 | Refuses to talk | 0.199 | 0.210 | 0.374 |
| **66** | **Repeats certain acts over and over; compulsions** | 0.167 | **0.508** | 0.203 |
| **68** | **Screams a lot** | **0.716** | -0.071 | 0.189 |
| 69 | Secretive, keeps things to self | 0.227 | 0.204 | 0.339 |
| **71** | **Self-conscious or easily embarrassed** | 0.009 | 0.228 | **0.583** |
| **72** | **Sets fires** | **0.556** | 0.123 | -0.184 |
| **74** | **Showing off or clowning** | **0.480** | 0.272 | -0.082 |
| **75** | **Too shy or timid** | -0.155 | 0.208 | **0.518** |
| **80** | **Stares blankly** | -0.058 | **0.741** | 0.161 |
| **81** | **Steals at home** | **0.690** | 0.146 | -0.123 |
| **82** | **Steals outside the home** | **0.660** | 0.207 | -0.162 |
| **84** | **Strange behavior** | 0.257 | **0.477** | 0.172 |
| **85** | **Strange ideas** | 0.169 | **0.462** | 0.203 |
| **86** | **Stubborn, sullen, or irritable** | **0.662** | -0.057 | 0.301 |
| **87** | **Sudden changes in mood or feelings** | **0.550** | 0.051 | 0.364 |
| **88** | **Sulks a lot** | **0.455** | 0.034 | **0.436** |
| **89** | **Suspicious** | **0.450** | 0.164 | 0.280 |
| **90** | **Swearing or obscene language** | **0.672** | -0.005 | 0.039 |
| 93 | Talks too much | 0.238 | 0.376 | 0.042 |
| **94** | **Teases a lot** | **0.649** | 0.036 | 0.052 |
| **95** | **Temper tantrums or hot temper** | **0.781** | -0.106 | 0.202 |
| **97** | **Threatens people** | **0.825** | -0.120 | 0.169 |
| **102** | **Underactive, slow moving, or lacks energy** | 0.054 | 0.293 | **0.424** |
| **103** | **Unhappy, sad, or depressed** | 0.280 | 0.101 | **0.560** |
| **106** | **Vandalism** | **0.729** | 0.215 | -0.145 |
| **109** | **Whining** | **0.420** | 0.066 | 0.295 |
| **111** | **Withdrawn, doesn't get involved with others** | 0.101 | 0.258 | **0.536** |
| **112** | **Worries** | 0.002 | 0.150 | **0.686** |
| **8,78** | **Inattentive*** | 0.103 | **0.869** | -0.079 |
| **20,21** | **Destroys things*** | **0.688** | 0.115 | 0.032 |

Latent factor correlations:
ADHD with
EXT 0.575
INT with
EXT 0.375
ADHD 0.344

Items in bold retained for the next round of exploratory analysis using data from the first random split half of the sample to specify which items loading on the specified factors in confirmatory factor analyses in the second random split half of the sample.

| Table S4. Initial exploratory correlated factors analysis in ESEM of CBCL items in the first half of the sample. | | | | | |
|---|---|---|---|---|---|
| Item | Brief wording | Ext | ADHD | Int | Somatic |
| **1** | **Acts too young for age** | 0.148 | **0.469** | 0.139 | -0.031 |
| **3** | **Argues a lot** | **0.761** | -0.023 | -0.028 | 0.180 |
| **4** | **Fails to finish things** | 0.183 | **0.574** | 0.071 | 0.034 |
| **5** | **There is very little enjoys** | 0.298 | 0.119 | **0.460** | -0.064 |
| 6 | Bowel movements outside toilet | 0.086 | 0.255 | 0.083 | -0.044 |
| **7** | **Bragging, boasting** | **0.486** | 0.137 | -0.175 | 0.257 |
| **9** | **Can't get mind off certain thoughts; obsessions** | 0.104 | **0.472** | 0.202 | 0.174 |
| **10** | **Can't sit still, restless, or hyperactive** | 0.210 | **0.732** | -0.186 | 0.069 |
| 11 | Clings to adults or too dependent | 0.099 | 0.308 | 0.274 | 0.166 |
| 12 | Complains of loneliness | 0.174 | 0.210 | 0.364 | 0.160 |
| **13** | **Confused or seems to be in a fog** | -0.094 | **0.647** | 0.306 | -0.033 |
| 14 | Cries a lot | 0.280 | 0.091 | 0.282 | 0.161 |
| **15** | **Cruel to animals** | **0.541** | -0.008 | 0.180 | -0.053 |
| **16** | **Cruelty, bullying, or meanness to others** | **0.850** | -0.114 | 0.067 | -0.020 |
| **17** | **Daydreams or gets lost in thoughts** | -0.152 | **0.678** | 0.160 | 0.069 |
| 18 | Deliberately harms self or attempts suicide | 0.352 | -0.056 | 0.362 | 0.108 |
| **19** | **Demands a lot of attention** | **0.451** | 0.260 | 0.003 | 0.252 |
| **22** | **Disobedient at home** | **0.835** | 0.035 | -0.083 | 0.061 |
| **23** | **Disobedient at school** | **0.672** | 0.322 | -0.153 | -0.153 |
| 24 | Doesn't eat well | 0.121 | 0.217 | 0.178 | 0.109 |
| **25** | **Doesn't get along with other kids** | **0.562** | 0.140 | 0.329 | -0.198 |
| **26** | **Doesn't seem to feel guilty after misbehaving** | **0.679** | 0.144 | -0.020 | -0.094 |
| **27** | **Easily jealous** | **0.522** | 0.011 | 0.164 | 0.161 |
| **28** | **Breaks rules at home, school or elsewhere** | **0.825** | 0.161 | -0.132 | -0.037 |
| 29 | Fears certain animals, situations, or places | -0.043 | 0.197 | 0.310 | 0.234 |
| **30** | **Fears going to school** | 0.103 | 0.036 | **0.545** | 0.111 |
| **31** | **Fears might think or do something bad** | 0.057 | 0.098 | **0.429** | 0.241 |
| **32** | **Feels has to be perfect** | -0.011 | -0.091 | **0.481** | 0.290 |
| **33** | **Feels or complains that no one loves them** | 0.429 | -0.077 | **0.450** | 0.137 |
| **34** | **Feels others are out to get** | 0.409 | 0.058 | **0.426** | 0.073 |
| **35** | **Feels worthless or inferior** | 0.196 | 0.076 | **0.575** | 0.152 |
| **36** | **Gets hurt a lot, accident prone** | 0.038 | **0.404** | -0.011 | 0.265 |
| **37** | **Gets in many fights** | **0.758** | -0.005 | 0.023 | -0.060 |
| 38 | Gets teased a lot | 0.214 | 0.290 | 0.345 | -0.051 |
| **39** | **Hangs around with others who get in trouble** | **0.496** | 0.197 | -0.058 | -0.048 |
| 40 | Hears sound or voices that aren't there | 0.113 | 0.279 | 0.179 | 0.127 |
| **41** | **Impulsive or acts without thinking** | **0.428** | **0.512** | -0.056 | 0.060 |
| **42** | **Would rather be alone than with others** | 0.053 | 0.151 | **0.589** | -0.148 |
| **43** | **Lying or cheating** | **0.614** | 0.218 | -0.090 | 0.026 |
| 44 | Bites fingernails | 0.106 | 0.201 | 0.024 | 0.096 |
| 45 | Nervous, highstrung, or tens | 0.042 | 0.291 | 0.390 | 0.287 |
| **46** | **Nervous movements or twitching** | 0.015 | **0.506** | 0.156 | 0.172 |
| 47 | Nightmares | 0.054 | 0.259 | 0.119 | 0.347 |

| | | | | | |
|---|---|---|---|---|---|
| 48 | Not liked by other kids | 0.371 | 0.244 | 0.393 | -0.150 |
| 49 | Constipated, doesn't move bowels | -0.004 | 0.147 | 0.181 | 0.306 |
| **50** | **Too fearful or anxious** | -0.118 | 0.252 | **0.525** | 0.330 |
| **51** | **Feels dizzy or lightheaded** | -0.025 | 0.124 | 0.165 | **0.522** |
| **52** | **Feels too guilty** | 0.010 | 0.053 | **0.507** | 0.325 |
| 54 | Overtired without good reason | 0.113 | 0.187 | 0.318 | 0.247 |
| **56A** | **Aches or pains** | 0.017 | 0.091 | -0.001 | **0.527** |
| **56B** | **Headaches** | 0.072 | 0.004 | 0.026 | **0.567** |
| **56C** | **Nausea, feels sick** | 0.020 | -0.023 | 0.010 | **0.840** |
| 56D | Problems with eyes | 0.002 | 0.052 | 0.181 | 0.261 |
| 56E | Rashes or other skin problems | 0.027 | 0.121 | 0.012 | 0.314 |
| **56F** | **Stomachaches** | 0.022 | -0.017 | 0.009 | **0.770** |
| **56G** | **Vomiting, throwing up** | -0.035 | 0.037 | -0.100 | **0.676** |
| **56H** | **Other physical problems** | 0.032 | 0.131 | 0.004 | **0.457** |
| **57** | **Physically attacks people** | **0.806** | -0.113 | 0.128 | -0.060 |
| 58 | Picks nose, skin, or other parts of body | 0.118 | 0.354 | 0.016 | 0.167 |
| 60 | Plays with own sex parts too much | 0.303 | 0.177 | -0.068 | 0.055 |
| **61** | **Poor school work** | 0.213 | **0.575** | 0.101 | -0.170 |
| **62** | **Poorly coordinated or clumsy** | -0.050 | **0.613** | 0.123 | 0.155 |
| 63 | Prefers being with older kids | 0.247 | 0.242 | 0.061 | 0.127 |
| 64 | Prefers being with younger kids | 0.029 | 0.372 | 0.232 | 0.065 |
| **65** | **Refuses to talk** | 0.208 | 0.109 | **0.559** | -0.130 |
| **66** | **Repeats certain acts over and over; compulsions** | 0.135 | **0.482** | 0.241 | 0.055 |
| **68** | **Screams a lot** | **0.703** | -0.065 | 0.097 | 0.169 |
| **69** | **Secretive, keeps things to self** | 0.224 | 0.128 | **0.455** | -0.039 |
| 70 | Sees things that aren't there | 0.159 | 0.259 | 0.149 | 0.151 |
| **71** | **Self-conscious or easily embarrassed** | 0.017 | 0.118 | **0.587** | 0.148 |
| **72** | **Sets fires** | **0.520** | 0.171 | 0.022 | -0.263 |
| **74** | **Showing off or clowning** | **0.465** | 0.324 | -0.216 | 0.141 |
| **75** | **Too shy or timid** | -0.162 | 0.089 | **0.677** | -0.033 |
| 76 | Sleeps less than most kids | -0.006 | 0.375 | 0.140 | 0.203 |
| 77 | Sleeps more than most kids | 0.081 | 0.155 | 0.305 | 0.048 |
| 79 | Speech problem | -0.032 | 0.386 | 0.124 | -0.102 |
| **80** | **Stares blankly** | -0.080 | **0.703** | 0.238 | -0.008 |
| **81** | **Steals at home** | **0.652** | 0.199 | -0.082 | -0.043 |
| **82** | **Steals outside the home** | **0.614** | 0.256 | -0.041 | -0.133 |
| 83 | Stores up too many things | 0.127 | 0.243 | 0.164 | 0.199 |
| **84** | **Strange behavior** | 0.224 | **0.457** | 0.266 | -0.025 |
| **85** | **Strange ideas** | 0.140 | **0.448** | 0.217 | 0.063 |
| **86** | **Stubborn, sullen, or irritable** | **0.666** | -0.083 | 0.181 | 0.192 |
| **87** | **Sudden changes in mood or feelings** | **0.550** | 0.008 | 0.284 | 0.180 |
| **88** | **Sulks a lot** | **0.466** | -0.040 | 0.378 | 0.171 |
| **89** | **Suspicious** | **0.439** | 0.116 | 0.298 | 0.072 |
| **90** | **Swearing or obscene language** | **0.655** | 0.013 | 0.042 | 0.014 |
| 91 | Talks about killing self | 0.264 | 0.065 | 0.359 | 0.215 |
| 92 | Talks or walks in sleep | 0.068 | 0.200 | -0.072 | 0.304 |

| | | | | | |
|---|---|---|---|---|---|
| **93** | **Talks too much** | 0.191 | **0.459** | -0.192 | 0.307 |
| **94** | **Teases a lot** | **0.638** | 0.062 | -0.036 | 0.109 |
| **95** | **Temper tantrums or hot temper** | **0.773** | -0.109 | 0.110 | 0.154 |
| **97** | **Threatens people** | **0.808** | -0.112 | 0.175 | 0.023 |
| 98 | Thumb-sucking | -0.016 | 0.103 | 0.069 | 0.128 |
| 100 | Trouble sleeping | 0.014 | 0.342 | 0.142 | 0.323 |
| **102** | **Underactive, slow moving, or lacks energy** | 0.042 | 0.231 | **0.454** | 0.092 |
| **103** | **Unhappy, sad, or depressed** | 0.292 | 0.010 | **0.516** | 0.183 |
| 104 | Unusually loud | 0.378 | 0.347 | -0.069 | 0.283 |
| **106** | **Vandalism** | **0.684** | 0.254 | -0.002 | -0.159 |
| 107 | Wets self during the day | 0.157 | 0.237 | 0.046 | 0.066 |
| 108 | Wets the bed | 0.169 | 0.163 | -0.034 | 0.004 |
| **109** | **Whining** | **0.413** | 0.064 | 0.103 | 0.287 |
| **111** | **Withdrawn, doesn't get involved with others** | 0.115 | 0.130 | **0.713** | -0.098 |
| **112** | **Worries** | -0.003 | 0.065 | **0.497** | 0.394 |
| **8,78** | **Inattentive*** | 0.085 | **0.865** | -0.042 | -0.022 |
| **20,21** | **Destroys things*** | **0.667** | 0.125 | 0.072 | -0.016 |
| 53,55 | Overeats/overweight* | 0.121 | 0.104 | 0.143 | 0.165 |

Factor correlations:
ADHD with
EXT 0.571
INT with
EXT 0.402
ADHD 0.378
SOMATIC with
EXT 0.244
ADHD 0.247
INT 0.360

*Composite item

Items in bold retained for the next round of exploratory analysis using data from the first random split half of the sample to specify which items loading on the specified factors in confirmatory factor analyses in the second random split half of the sample.

Table S5. Standardized factor loadings from the confirmatory bifactor model plus two specific factors based on CBCL items in the second split half of the wave 1 ABCD Study (N = 5934).

| Item | Brief wording | General | Externalizing | Internalizing |
|---|---|---|---|---|
| 1 | Acts too young for age | **0.518** | 0.274 | |
| 3 | Argues a lot | **0.647** | **0.428** | |
| 4 | Fails to finish things | **0.595** | **0.411** | |
| 5 | There is very little enjoys | **0.725** | | -0.047[a] |
| 7 | Bragging, boasting | 0.357 | **0.404** | |
| 9 | Can't get mind off certain thoughts; obsessions | **0.671** | 0.210 | |
| 10 | Can't sit still, restless, or hyperactive | **0.505** | **0.544** | |
| 15 | Cruel to animals | **0.462** | **0.493** | |
| 16 | Cruelty, bullying, or meanness to others | **0.530** | **0.516** | |
| 19 | Demands a lot of attention | **0.634** | 0.343 | |
| 22 | Disobedient at home | **0.595** | **0.597** | |
| 23 | Disobedient at school | **0.435** | **0.682** | |
| 25 | Doesn't get along with other kids | **0.642** | 0.346 | |
| 26 | Doesn't seem to feel guilty after misbehaving | **0.542** | **0.517** | |
| 27 | Easily jealous | **0.645** | 0.247 | |
| 28 | Breaks rules at home, school or elsewhere | **0.526** | **0.714** | |
| 30 | Fears going to school | **0.620** | | 0.289 |
| 31 | Fears might think or do something bad | **0.559** | | 0.300 |
| 32 | Feels has to be perfect | 0.361 | | 0.369 |
| 33 | Feels or complains that no one loves them | **0.803** | | 0.002[a] |
| 34 | Feels others are out to get | **0.789** | | -0.054[a] |
| 35 | Feels worthless or inferior | **0.732** | | 0.196 |
| 37 | Gets in many fights | **0.530** | **0.537** | |
| 39 | Hangs around with others who get in trouble | **0.359** | **0.508** | |
| 41 | Impulsive or acts without thinking | **0.605** | **0.524** | |
| 42 | Would rather be alone than with others | **0.583** | | 0.095[d] |
| 43 | Lying or cheating | **0.460** | **0.605** | |
| 50 | Too fearful or anxious | **0.615** | | 0.397 |
| 51 | Feels dizzy or lightheaded | **0.467** | | **0.475** |
| 52 | Feels too guilty | **0.574** | | **0.451** |
| 56A | Aches or pains | **0.417** | | **0.411** |
| 56B | Headaches | 0.342 | | **0.456** |
| 56C | Nausea, feels sick | **0.435** | | **0.736** |
| 56F | Stomachaches | 0.367 | | **0.678** |
| 56G | Vomiting, throwing up | 0.297 | | **0.511** |
| 56H | Other physical problems | **0.454** | | 0.371 |
| 57 | Physically attacks people | **0.592** | **0.451** | |
| 61 | Poor school work | **0.507** | **0.425** | |
| 65 | Refuses to talk | **0.686** | | 0.052[a] |
| 66 | Repeats certain acts over and over; compulsions | **0.631** | 0.242 | |
| 68 | Screams a lot | **0.645** | 0.350 | |
| 69 | Secretive, keeps things to self | **0.703** | | 0.025[a] |

| | | | | |
|---|---|---|---|---|
| 71 | Self-conscious or easily embarrassed | **0.593** | | 0.341 |
| 72 | Sets fires | 0.328 | **0.503** | |
| 74 | Showing off or clowning | 0.392 | **0.493** | |
| 75 | Too shy or timid | **0.442** | | 0.302 |
| 80 | Stares blankly | **0.645** | 0.131 | |
| 81 | Steals at home | **0.431** | **0.704** | |
| 82 | Steals outside the home | 0.388 | **0.721** | |
| 84 | Strange behavior | **0.732** | 0.283 | |
| 85 | Strange ideas | **0.653** | 0.211 | |
| 86 | Stubborn, sullen, or irritable | **0.735** | 0.265 | |
| 87 | Sudden changes in mood or feelings | **0.806** | 0.164 | |
| 88 | Sulks a lot | **0.784** | | $0.071^{c}$ |
| 89 | Suspicious | **0.701** | 0.259 | |
| 90 | Swearing or obscene language | **0.485** | **0.440** | |
| 93 | Talks too much | **0.432** | 0.351 | |
| 94 | Teases a lot | **0.513** | **0.467** | |
| 95 | Temper tantrums or hot temper | **0.680** | 0.350 | |
| 97 | Threatens people | **0.620** | **0.554** | |
| 102 | Underactive, slow moving, or lacks energy | **0.606** | | 0.232 |
| 103 | Unhappy, sad, or depressed | **0.771** | | 0.192 |
| 106 | Vandalism | **0.544** | **0.573** | |
| 111 | Withdrawn, doesn't get involved with others | **0.708** | | 0.160 |
| 112 | Worries | **0.596** | | **0.441** |
| 8,78 | Inattentive* | **0.580** | **0.522** | |
| 20,21 | Destroys things belonging to self or others* | **0.616** | **0.515** | |

*Composite of two nearly synonymous and highly correlated items.
$^{a}p > .05$; $^{b}p < .05$; $^{c}p < .01$; $^{d}p < .001$; All loadings on all factors without a superscript $p < .0001$.

Ext = externalizing; Int = internalizing; ADHD = attention deficit hyperactivity disorder/sluggish cognitive tempo; soma = somatization.

Table S6. Standardized factor loadings from the confirmatory bifactor model plus three specific factors based on CBCL items in the second split half of the wave 1 ABCD Study (N = 5934).

| Item | Brief wording | General | Conduct | ADHD | Int |
|---|---|---|---|---|---|
| 1 | Acts too young for age | **0.572** | | 0.265 | |
| 3 | Argues a lot | **0.743** | 0.256 | | |
| 4 | Fails to finish things | **0.694** | | 0.329 | |
| 7 | Bragging, boasting | **0.468** | 0.264 | | |
| 9 | Can't get mind off certain thoughts; obsessions | **0.686** | | 0.232 | |
| 10 | Can't sit still, restless, or hyperactive | **0.649** | | **0.460** | |
| 13 | Confused or seems to be in a fog | **0.599** | | **0.505** | |
| 15 | Cruel to animals | **0.552** | **0.410** | | |
| 16 | Cruelty, bullying, or meanness to others | **0.610** | **0.477** | | |
| 17 | Daydreams or gets lost in thoughts | **0.509** | | **0.474** | |
| 19 | Demands a lot of attention | **0.727** | 0.103 | | |
| 22 | Disobedient at home | **0.729** | **0.458** | | |
| 23 | Disobedient at school | **0.630** | **0.458** | | |
| 25 | Doesn't get along with other kids | **0.698** | 0.223 | | |
| 26 | Doesn't seem to feel guilty after misbehaving | **0.651** | 0.388 | | |
| 27 | Easily jealous | **0.679** | 0.134 | | |
| 28 | Breaks rules at home, school or elsewhere | **0.701** | **0.557** | | |
| 30 | Fears going to school | **0.554** | | | 0.397 |
| 31 | Fears might think or do something bad | **0.495** | | | **0.412** |
| 32 | Feels has to be perfect | 0.299 | | | **0.465** |
| 33 | Feels or complains that no one loves them | **0.726** | | | 0.193 |
| 34 | Feels others are out to get | **0.737** | -0.016[a] | | |
| 35 | Feels worthless or inferior | **0.662** | | | 0.364 |
| 37 | Gets in many fights | **0.628** | **0.455** | | |
| 39 | Hangs around with others who get in trouble | **0.484** | 0.373 | | |
| 41 | Impulsive or acts without thinking | **0.794** | | 0.186 | |
| 43 | Lying or cheating | **0.600** | **0.484** | | |
| 46 | Nervous movements or twitching | **0.540** | | 0.323 | |
| 50 | Too fearful or anxious | **0.557** | | | **0.485** |
| 51 | Feels dizzy or lightheaded | **0.433** | | | **0.488** |
| 52 | Feels too guilty | **0.513** | | | **0.538** |
| 56A | Aches or pains | 0.383 | | | **0.404** |
| 56B | Headaches | 0.313 | | | **0.442** |
| 56C | Nausea, feels sick | **0.407** | | | **0.707** |
| 56F | Stomachaches | 0.342 | | | **0.643** |
| 56G | Vomiting, throwing up | 0.282 | | | **0.464** |
| 57 | Physically attacks people | **0.653** | 0.413 | | |
| 61 | Poor school work | **0.616** | | 0.346 | |
| 62 | Poorly coordinated or clumsy | **0.572** | | 0.288 | |
| 66 | Repeats certain acts over and over; compulsions | **0.656** | | 0.301 | |
| 68 | Screams a lot | **0.703** | 0.234 | | |
| 71 | Self-conscious or easily embarrassed | **0.517** | | | **0.466** |
| 72 | Sets fires | **0.455** | 0.387[b] | | |
| 74 | Showing off or clowning | **0.556** | 0.219 | | |
| 75 | Too shy or timid | 0.359 | | | **0.417** |

| | | | | | |
|---|---|---|---|---|---|
| 80 | Stares blankly | **0.600** | | **0.477** | |
| 81 | Steals at home | **0.550** | **0.676** | | |
| 82 | Steals outside the home | **0.520** | **0.675** | | |
| 84 | Strange behavior | **0.778** | | 0.215 | |
| 85 | Strange ideas | **0.673** | | 0.203 | |
| 86 | Stubborn, sullen, or irritable | **0.778** | 0.114 | | |
| 87 | Sudden changes in mood or feelings | **0.831** | -0.025[a] | | |
| 88 | Sulks a lot | **0.775** | -0.126 | | |
| 89 | Suspicious | **0.721** | 0.162 | | |
| 90 | Swearing or obscene language | **0.546** | **0.417** | | |
| 94 | Teases a lot | **0.604** | 0.362 | | |
| 95 | Temper tantrums or hot temper | **0.740** | 0.227 | | |
| 97 | Threatens people | **0.698** | **0.507** | | |
| 102 | Underactive, slow moving, or lacks energy | **0.553** | | | 0.318 |
| 103 | Unhappy, sad, or depressed | **0.697** | | | 0.337 |
| 106 | Vandalism | **0.640** | **0.498** | | |
| 109 | Whining | **0.662** | -0.024[a] | | |
| 111 | Withdrawn, doesn't get involved with others | **0.618** | | | 0.294 |
| 112 | Worries | **0.535** | | | **0.534** |
| 8,78 | Inattentive* | **0.680** | | **0.636** | |
| 20,21 | Destroys things belonging to self or others* | **0.709** | **0.408** | | |

*Composite of two nearly synonymous and highly correlated items.
[a]$p > .05$; [b]$p < .005$. All loadings without a superscript, $p < .0001$.

Int = internalizing; ADHD = attention deficit hyperactivity disorder/sluggish cognitive tempo.
Standardized loadings ≥ 0.400 in bold.

Table S7. Standardized factor loadings from the confirmatory second-order model based on three correlated factors of CBCL items in the second split half of the wave 1 ABCD Study (N = 5934).

| Loadings of CBCL Items on Lower-Order Factors | | | | |
|---|---|---|---|---|
| Item | Brief wording | Conduct | ADHD | Internalizing |
| 1 | Acts too young for age | | 0.643 | |
| 3 | Argues a lot | 0.792 | | |
| 4 | Fails to finish things | | 0.785 | |
| 7 | Bragging, boasting | 0.526 | | |
| 9 | Can't get mind off certain thoughts; obsessions | | 0.755 | |
| 10 | Can't sit still, restless, or hyperactive | | 0.769 | |
| 13 | Confused or seems to be in a fog | | 0.723 | |
| 15 | Cruel to animals | 0.654 | | |
| 16 | Cruelty, bullying, or meanness to others | 0.729 | | |
| 17 | Daydreams or gets lost in thoughts | | 0.625 | |
| 19 | Demands a lot of attention | 0.739 | | |
| 22 | Disobedient at home | 0.836 | | |
| 23 | Disobedient at school | 0.743 | | |
| 25 | Doesn't get along with other kids | 0.739 | | |
| 26 | Doesn't seem to feel guilty after misbehaving | 0.741 | | |
| 27 | Easily jealous | 0.699 | | |
| 28 | Breaks rules at home, school or elsewhere | 0.845 | | |
| 30 | Fears going to school | | | 0.706 |
| 31 | Fears might think or do something bad | | | 0.649 |
| 32 | Feels has to be perfect | | | 0.457 |
| 33 | Feels or complains that no one loves them | | | 0.850 |
| 34 | Feels others are out to get | 0.721 | | |
| 35 | Feels worthless or inferior | | | 0.812 |
| 37 | Gets in many fights | 0.741 | | |
| 39 | Hangs around with others who get in trouble | 0.573 | | |
| 41 | Impulsive or acts without thinking | | 0.858 | |
| 43 | Lying or cheating | 0.720 | | |
| 46 | Nervous movements or twitching | | 0.623 | |
| 50 | Too fearful or anxious | | | 0.738 |
| 51 | Feels dizzy or lightheaded | | | 0.607 |
| 52 | Feels too guilty | | | 0.705 |
| 56A | Aches or pains | | | 0.526 |
| 56B | Headaches | | | 0.464 |
| 56C | Nausea, feels sick | | | 0.656 |
| 56F | Stomachaches | | | 0.569 |
| 56G | Vomiting, throwing up | | | 0.439 |
| 57 | Physically attacks people | 0.751 | | |
| 61 | Poor school work | | 0.706 | |
| 62 | Poorly coordinated or clumsy | | 0.647 | |
| 66 | Repeats certain acts over and over; compulsions | | 0.736 | |
| 68 | Screams a lot | 0.747 | | |
| 71 | Self-conscious or easily embarrassed | | | 0.688 |
| 72 | Sets fires | 0.550 | | |
| 74 | Showing off or clowning | 0.600 | | |

| | | | | |
|---|---|---|---|---|
| 75 | Too shy or timid | | | 0.508 |
| 80 | Stares blankly | | 0.720 | |
| 81 | Steals at home | 0.754 | | |
| 82 | Steals outside the home | 0.734 | | |
| 84 | Strange behavior | | 0.847 | |
| 85 | Strange ideas | | 0.734 | |
| 86 | Stubborn, sullen, or irritable | 0.790 | | |
| 87 | Sudden changes in mood or feelings | 0.812 | | |
| 88 | Sulks a lot | 0.734 | | |
| 89 | Suspicious | 0.744 | | |
| 90 | Swearing or obscene language | 0.647 | | |
| 94 | Teases a lot | 0.687 | | |
| 95 | Temper tantrums or hot temper | 0.781 | | |
| 97 | Threatens people | 0.827 | | |
| 102 | Underactive, slow moving, or lacks energy | | | 0.686 |
| 103 | Unhappy, sad, or depressed | | | 0.848 |
| 106 | Vandalism | 0.771 | | |
| 109 | Whining | 0.646 | | |
| 111 | Withdrawn, doesn't get involved with others | | | 0.752 |
| 112 | Worries | | | 0.729 |
| 8,78 | Inattentive* | | 0.834 | |
| 20,21 | Destroys things belonging to self or others* | 0.804 | | |
| Loadings of Lower-Order Factors on Second-Order General Factor | | | | |
| General Factor | | 0.892 | 0.894 | 0.761 |

*Composite of two nearly synonymous and highly correlated items.
All loadings, $p < .0001$.
Int = internalizing; ADHD = attention deficit hyperactivity disorder/sluggish cognitive tempo.

Table S8. Standardized factor loadings from the confirmatory bifactor model plus four specific factors based on CBCL items in the second split half of the wave 1 ABCD Study (N = 5934).

| Item | Brief wording | General | Ext | ADHD | Int | Somatic |
|---|---|---|---|---|---|---|
| 1 | Acts too young for age | **0.573** | | 0.259 | | |
| 3 | Argues a lot | **0.719** | 0.315 | | | |
| 4 | Fails to finish things | **0.695** | | 0.323 | | |
| 5 | There is very little enjoys | **0.657** | | | 0.130 | |
| 7 | Bragging, boasting | **0.452** | 0.293 | | | |
| 9 | Can't get mind off certain thoughts; obsessions | **0.689** | | 0.225 | | |
| 10 | Can't sit still, restless, or hyperactive | **0.644** | | **0.476** | | |
| 13 | Confused or seems to be in a fog | **0.617** | | **0.462** | | |
| 15 | Cruel to animals | **0.550** | **0.406** | | | |
| 16 | Cruelty, bullying, or meanness to others | **0.592** | **0.501** | | | |
| 17 | Daydreams or gets lost in thoughts | **0.519** | | **0.451** | | |
| 19 | Demands a lot of attention | **0.716** | 0.143 | | | |
| 22 | Disobedient at home | **0.704** | **0.501** | | | |
| 23 | Disobedient at school | **0.617** | **0.467** | | | |
| 25 | Doesn't get along with other kids | **0.697** | 0.229 | | | |
| 26 | Doesn't seem to feel guilty after misbehaving | **0.644** | 0.398 | | | |
| 27 | Easily jealous | **0.669** | 0.168 | | | |
| 28 | Breaks rules at home, school or elsewhere | **0.682** | **0.578** | | | |
| 30 | Fears going to school | **0.574** | | | 0.363 | |
| 31 | Fears might think or do something bad | **0.504** | | | **0.444** | |
| 32 | Feels has to be perfect | 0.307 | | | **0.523** | |
| 33 | Feels or complains that no one loves them | **0.749** | 0.019[a] | | | |
| 34 | Feels others are out to get | **0.730** | 0.015[a] | | | |
| 35 | Feels worthless or inferior | **0.674** | | | 0.335 | |
| 36 | Gets hurt a lot, accident prone | **0.492** | | 0.245 | | |
| 37 | Gets in many fights | **0.621** | **0.460** | | | |
| 39 | Hangs around with others who get in trouble | **0.482** | 0.365 | | | |
| 41 | Impulsive or acts without thinking | **0.790** | | 0.191 | | |
| 42 | Would rather be alone than with others | **0.509** | | | 0.357 | |
| 43 | Lying or cheating | **0.593** | **0.485** | | | |
| 46 | Nervous movements or twitching | **0.546** | | 0.310 | | |
| 50 | Too fearful or anxious | **0.566** | | | **0.510** | |
| 51 | Feels dizzy or lightheaded | **0.497** | | | | **0.456** |
| 52 | Feels too guilty | **0.533** | | | **0.520** | |
| 56A | Aches or pains | **0.433** | | | | **0.454** |
| 56B | Headaches | 0.362 | | | | **0.520** |
| 56C | Nausea, feels sick | **0.481** | | | | **0.762** |
| 56F | Stomachaches | **0.409** | | | | **0.729** |
| 56G | Vomiting, throwing up | 0.322 | | | | **0.602** |
| 56H | Other physical problems | **0.461** | | | | **0.422** |
| 57 | Physically attacks people | **0.633** | **0.448** | | | |
| 61 | Poor school work | **0.617** | | 0.336 | | |

| | | | | | |
|---|---|---|---|---|---|
| 62 | Poorly coordinated or clumsy | **0.582** | | 0.355 | |
| 65 | Refuses to talk | **0.608** | | | 0.298 |
| 66 | Repeats certain acts over and over; compulsions | **0.665** | | 0.279 | |
| 68 | Screams a lot | **0.693** | 0.268 | | |
| 69 | Secretive, keeps things to self | **0.629** | | | 0.245 |
| 71 | Self-conscious or easily embarrassed | **0.523** | | | **0.556** |
| 72 | Sets fires | **0.439** | 0.397[c] | | |
| 74 | Showing off or clowning | **0.544** | 0.240 | | |
| 75 | Too shy or timid | 0.362 | | | **0.567** |
| 80 | Stares blankly | **0.616** | | **0.435** | |
| 81 | Steals at home | **0.548** | **0.663** | | |
| 82 | Steals outside the home | **0.518** | **0.663** | | |
| 84 | Strange behavior | **0.788** | | 0.184 | |
| 85 | Strange ideas | **0.684** | | 0.185 | |
| 86 | Stubborn, sullen, or irritable | **0.759** | 0.176 | | |
| 87 | Sudden changes in mood or feelings | **0.817** | 0.035[a] | | |
| 88 | Sulks a lot | **0.766** | -0.071[b] | | |
| 89 | Suspicious | **0.724** | 0.169 | | |
| 90 | Swearing or obscene language | **0.534** | **0.432** | | |
| 93 | Talks too much | **0.542** | | 0.200 | |
| 94 | Teases a lot | **0.595** | 0.377 | | |
| 95 | Temper tantrums or hot temper | **0.717** | 0.287 | | |
| 97 | Threatens people | **0.680** | **0.532** | | |
| 102 | Underactive, slow moving, or lacks energy | **0.580** | | | 0.264 |
| 103 | Unhappy, sad, or depressed | **0.714** | | | 0.303 |
| 106 | Vandalism | **0.634** | **0.500** | | |
| 109 | Whining | **0.646** | 0.033[a] | | |
| 111 | Withdrawn, doesn't get involved with others | **0.628** | | | **0.439** |
| 112 | Worries | **0.548** | | | **0.544** |
| 8,78 | Inattentive* | **0.679** | | **0.634** | |
| 20,21 | Destroys things* | **0.701** | **0.419** | | |

*Composite of two nearly synonymous and highly correlated items.
[a]$p > .05$; [b]$p < .05$; [c]$p < .01$; [d]$p < .001$; All loadings on all factors without a superscript $p < .0001$.

Ext = externalizing; Int = internalizing; ADHD = attention deficit hyperactivity disorder/sluggish cognitive tempo.
Standardized loadings ≥ 0.400 in bold.

Table S9. Standardized factor loadings from the confirmatory second-order model based on four correlated factors of CBCL items in the second split half of the wave 1 ABCD Study (N = 5934).

| Loadings of CBCL Items on Lower-Order Factors | | | | | |
|---|---|---|---|---|---|
| Item | Brief wording | Ext | ADHD | Int | Somatic |
| 1 | Acts too young for age | | 0.640 | | |
| 3 | Argues a lot | 0.787 | | | |
| 4 | Fails to finish things | | 0.781 | | |
| 5 | There is very little enjoys | | | 0.745 | |
| 7 | Bragging, boasting | 0.521 | | | |
| 9 | Can't get mind off certain thoughts; obsessions | | 0.753 | | |
| 10 | Can't sit still, restless, or hyperactive | | 0.764 | | |
| 13 | Confused or seems to be in a fog | | 0.727 | | |
| 15 | Cruel to animals | 0.655 | | | |
| 16 | Cruelty, bullying, or meanness to others | 0.724 | | | |
| 17 | Daydreams or gets lost in thoughts | | 0.627 | | |
| 19 | Demands a lot of attention | 0.740 | | | |
| 22 | Disobedient at home | 0.831 | | | |
| 23 | Disobedient at school | 0.738 | | | |
| 25 | Doesn't get along with other kids | 0.742 | | | |
| 26 | Doesn't seem to feel guilty after misbehaving | 0.742 | | | |
| 27 | Easily jealous | 0.700 | | | |
| 28 | Breaks rules at home, school or elsewhere | 0.839 | | | |
| 30 | Fears going to school | | | 0.697 | |
| 31 | Fears might think or do something bad | | | 0.641 | |
| 32 | Feels has to be perfect | | | 0.445 | |
| 33 | Feels or complains that no one loves them | 0.743 | | | |
| 34 | Feels others are out to get | 0.723 | | | |
| 35 | Feels worthless or inferior | | | 0.801 | |
| 36 | Gets hurt a lot, accident prone | | 0.554 | | |
| 37 | Gets in many fights | 0.740 | | | |
| 39 | Hangs around with others who get in trouble | 0.572 | | | |
| 41 | Impulsive or acts without thinking | | 0.851 | | |
| 42 | Would rather be alone than with others | | | 0.628 | |
| 43 | Lying or cheating | 0.718 | | | |
| 46 | Nervous movements or twitching | | 0.623 | | |
| 50 | Too fearful or anxious | | | 0.724 | |
| 51 | Feels dizzy or lightheaded | | | | 0.803 |
| 52 | Feels too guilty | | | 0.690 | |
| 56A | Aches or pains | | | | 0.715 |
| 56B | Headaches | | | | 0.634 |
| 56C | Nausea, feels sick | | | | 0.870 |
| 56F | Stomachaches | | | | 0.758 |
| 56G | Vomiting, throwing up | | | | 0.606 |
| 56H | Other physical problems | | | | 0.741 |
| 57 | Physically attacks people | 0.747 | | | |

| | | | | | |
|---|---|---|---|---|---|
| 61 | Poor school work | | 0.702 | | |
| 62 | Poorly coordinated or clumsy | | 0.668 | | |
| 65 | Refuses to talk | | | 0.722 | |
| 66 | Repeats certain acts over and over; compulsions | | 0.738 | | |
| 68 | Screams a lot | 0.750 | | | |
| 69 | Secretive, keeps things to self | | | 0.736 | |
| 71 | Self-conscious or easily embarrassed | | | 0.684 | |
| 72 | Sets fires | 0.541 | | | |
| 74 | Showing off or clowning | 0.596 | | | |
| 75 | Too shy or timid | | | 0.522 | |
| 80 | Stares blankly | | 0.722 | | |
| 81 | Steals at home | 0.751 | | | |
| 82 | Steals outside the home | 0.729 | | | |
| 84 | Strange behavior | | 0.848 | | |
| 85 | Strange ideas | | 0.740 | | |
| 86 | Stubborn, sullen, or irritable | 0.789 | | | |
| 87 | Sudden changes in mood or feelings | 0.813 | | | |
| 88 | Sulks a lot | 0.738 | | | |
| 89 | Suspicious | 0.752 | | | |
| 90 | Swearing or obscene language | 0.644 | | | |
| 93 | Talks too much | | 0.597 | | |
| 94 | Teases a lot | 0.687 | | | |
| 95 | Temper tantrums or hot temper | 0.778 | | | |
| 97 | Threatens people | 0.823 | | | |
| 102 | Underactive, slow moving, or lacks energy | | | 0.684 | |
| 103 | Unhappy, sad, or depressed | | | 0.840 | |
| 106 | Vandalism | 0.770 | | | |
| 109 | Whining | 0.645 | | | |
| 111 | Withdrawn, doesn't get involved with others | | | 0.773 | |
| 112 | Worries | | | 0.712 | |
| 8,78 | Inattentive* | | 0.828 | | |
| 20,21 | Destroys things* | 0.803 | | | |
| Loadings of Lower-Order Factors on Second-Order General Factor | | | | | |
| General Factor | | 0.882 | 0.897 | 0.813 | 0.575 |

*Composite of two nearly synonymous and highly correlated items.
All loadings, $p < .0001$.
Ext = externalizing; Int = internalizing; ADHD = attention deficit hyperactivity disorder/sluggish cognitive tempo.

Table S10. Fit statistics for alternative confirmatory models of the final CBCL items based on the second split half of ABCD Study wave 1 sample.

Bifactor+Three Specific Factors

Chi-Square Test of Model Fit = 8757.891*; Degrees of Freedom = 2013

| | |
|---|---|
| RMSEA (90% CI) | 0.024 (0.023-0.024) |
| CFI | 0.943 |
| TLI | 0.940 |
| SRMR | 0.063 |

Bifactor+Four Specific Factors

Chi-Square Test of Model Fit = 9641.816*; Degrees of Freedom = 2482

| | |
|---|---|
| RMSEA (90% CI) | 0.022 (0.022- 0.023) |
| CFI | 0.943 |
| TLI | 0.939 |
| SRMR | 0.063 |

Second-Order Based on Three Correlated Lower-Order Factors

Chi-Square Test of Model Fit = 10639.825*; Degrees of Freedom = 2076

| | |
|---|---|
| RMSEA (90% CI) | 0.026 (0.026-0.027) |
| CFI | 0.928 |
| TLI | 0.926 |
| SRMR | 0.075 |

Second-Order Based on Four Lower-Order Correlated Factors

Chi-Square Test of Model Fit = 11306.504*; Degrees of Freedom = 2551

| | |
|---|---|
| RMSEA (90% CI) | 0.024 (0.024-0.025) |
| CFI | 0.930 |
| TLI | 0.928 |
| SRMR | 0.074 |

* P < .0001

RMSEA (90% CI) = root mean square error of approximation; CFI = comparative fit index; TLI = Tucker-Lewis index; SRMR = standardized root mean square residual.

Table S11. Results of simultaneous regressions of each independently measured criterion variable on the latent general factor and four specific factors defined in bifactor models and on demographic covariates of no interest (age, sex, and race-ethnicity) in the second split half of the ABCD Study sample (N = 5926).

| Bifactor Model with Four Specific Factors | | | | | | | | | | |
|---|---|---|---|---|---|---|---|---|---|---|
| | General | | Internalizing | | Externalizing | | ADHD | | Somatic | |
| | B | P < | β | P < | β | P < | β | P < | β | P < |
| *Functional Impairment (Binary)* | | | | | | | | | | |
| Detention/suspension | **0.379** | **0.001** | **-0.142** | **0.001** | **0.402** | **0.001** | 0.036 | 0.341 | -0.028 | 0.472 |
| Mental health service | **0.571** | **0.001** | **0.123** | **0.001** | **0.121** | **0.001** | 0.066 | 0.035 | 0.030 | 0.299 |
| Special Education (Behavior) | **0.281** | **0.001** | 0.058 | 0.368 | **0.271** | **0.001** | 0.110 | 0.042 | -0.029 | 0.653 |
| Special Education (Learning) | **0.298** | **0.001** | 0.019 | 0.774 | 0.047 | 0.305 | **0.165** | **0.011** | -0.029 | 0.633 |
| *Youth-Reported Harmful Behaviors (Binary)* | | | | | | | | | | |
| Suicidal Behavior | **0.274** | **0.001** | 0.038 | 0.411 | 0.092 | 0.080 | -0.001 | 0.988 | -0.021 | 0.704 |
| Nonsuicidal self-harm | **0.221** | **0.001** | **0.082** | **0.025** | 0.078 | 0.078 | 0.085 | 0.048 | -0.029 | 0.461 |
| *Dispositional Constructs (Standardized Continuous)* | | | | | | | | | | |
| UPPS Low premeditation | **0.129** | **0.001** | **-0.066** | **0.001** | **0.101** | **0.001** | **0.113** | **0.001** | **-0.094** | **0.001** |
| UPPS Low perseverance | **0.128** | **0.001** | **0.043** | **0.022** | **0.073** | **0.001** | **0.237** | **0.001** | -0.024 | 0.265 |
| UPPS Negative urgency | **0.165** | **0.001** | **-0.044** | **0.023** | **0.098** | **0.001** | 0.002 | 0.933 | **-0.078** | **0.001** |
| UPPS Positive urgency | **0.113** | **0.001** | **-0.100** | **0.001** | **0.094** | **0.001** | **0.081** | **0.001** | **-0.078** | **0.001** |
| UPPS Sensation seeking | 0.001 | 0.945 | **-0.096** | **0.001** | **0.090** | **0.001** | 0.032 | 0.192 | -0.006 | 0.804 |
| Prosociality | **-0.064** | **0.001** | -0.029 | 0.157 | **-0.077** | **0.001** | -0.031 | 0.202 | **0.061** | **0.005** |
| *Latent General Executive Functioning Test Scores* | | | | | | | | | | |
| Executive functioning | **-0.096** | **0.001** | 0.030 | 0.141 | **-0.223** | **0.001** | **-0.318** | **0.001** | 0.023 | 0.302 |

Coefficients in bold are significant after FDR correction (adopting a 5% false discovery rate) for 65 tests.
ADHD = attention-deficit hyperactivity disorder/sluggish cognitive tempo
Continuous dispositional and executive functioning measures were normalized to a mean of 0 and standard deviation of 1.

Table S12. Results of regressions of each independently measured criterion variable on the latent general factor and demographic covariates of no interest (age, sex, and race-ethnicity) and in separate models simultaneously on four lower-order factors defined in second-order factor models and on demographic covariates of no interest (age, sex, and race-ethnicity) in the second split half of the ABCD Study sample (N = 5926).

| Second-Order Model Based on Four Lower-Order Factors | | | | | | | | | | |
|---|---|---|---|---|---|---|---|---|---|---|
| Criterion Variable | Factors | | | | | | | | | |
| | General | | Internalizing | | Externalizing | | ADHD | | Somatic | |
| | β | P < | β | P < | β | P < | β | P < | β | P < |
| Regressions of Criterion Variables on Only the Second-Order General Factor and Covariates | | | | | | | | | | |
| *Functional Impairment (Binary)* | | | | | | | | | | |
| Detention/suspension | **0.485** | **0.001** | | | | | | | | |
| Mental health service | **0.629** | **0.001** | | | | | | | | |
| Special education (Behavior) | **0.375** | **0.001** | | | | | | | | |
| Special education (Learning) | **0.335** | **0.001** | | | | | | | | |
| *Youth-Reported Harmful Behaviors (Binary)* | | | | | | | | | | |
| Suicidal behavior | **0.301** | **0.001** | | | | | | | | |
| Nonsuicidal self-harm | **0.263** | **0.001** | | | | | | | | |
| *Youth-Rated Dispositional Constructs (Standardized Continuous)* | | | | | | | | | | |
| UPPS low premeditation | **0.157** | **0.001** | | | | | | | | |
| UPPS low perseverance | **0.186** | **0.001** | | | | | | | | |
| UPPS negative urgency | **0.180** | **0.001** | | | | | | | | |
| UPPS positive urgency | **0.130** | **0.001** | | | | | | | | |
| UPPS sensation seeking | 0.014 | 0.440 | | | | | | | | |
| Prosociality | **-0.087** | **0.001** | | | | | | | | |
| *Latent General Executive Functioning Test Scores* | | | | | | | | | | |
| Executive functioning | **-0.192** | **0.001** | | | | | | | | |
| Simultaneous Regressions of Criterion Variables on Only the Four Lower-Order Factors and Covariates | | | | | | | | | | |
| *Functional Impairment (Binary)* | | | | | | | | | | |
| Detention/suspension | | | **-0.289** | **0.001** | **0.708** | **0.001** | 0.047 | 0.407 | -0.045 | 0.316 |
| Mental health service | | | **0.230** | **0.001** | **0.259** | **0.001** | **0.141** | **0.003** | 0.042 | 0.214 |
| Special education (Behavior) | | | -0.001 | 0.991 | **0.388** | **0.001** | 0.034 | 0.691 | -0.075 | 0.276 |
| Special education (Learning) | | | 0.020 | 0.827 | 0.049 | 0.586 | **0.307** | **0.001** | -0.043 | 0.535 |
| *Youth-Reported Harmful Behaviors (Binary)* | | | | | | | | | | |
| Suicidal behavior | | | 0.096 | 0.196 | 0.096 | 0.041 | 0.068 | 0.415 | -0.030 | 0.642 |
| Nonsuicidal self-harm | | | 0.125 | 0.032 | 0.077 | 0.265 | 0.110 | 0.088 | -0.051 | 0.282 |
| *Youth-Rated Dispositional Constructs (Standardized Continuous)* | | | | | | | | | | |
| UPPS low premeditation | | | **-0.129** | **0.001** | **0.125** | **0.001** | **0.235** | **0.001** | **-0.123** | **0.001** |
| UPPS low perseverance | | | -0.010 | 0.736 | -0.068 | 0.038 | **0.325** | **0.001** | **-0.076** | **0.002** |
| UPPS negative urgency | | | -0.043 | 0.154 | **0.232** | **0.001** | 0.047 | 0.191 | **-0.093** | **0.001** |
| UPPS positive urgency | | | **-0.173** | **0.001** | **0.168** | **0.001** | **0.189** | **0.001** | **-0.101** | **0.001** |
| UPPS sensation seeking | | | **-0.176** | **0.001** | **0.121** | **0.001** | 0.063 | 0.087 | -0.016 | 0.556 |
| Prosociality | | | -0.036 | 0.261 | **-0.099** | **0.004** | -0.010 | 0.776 | **0.083** | **0.001** |
| *Latent General Executive Functioning Test Scores* | | | | | | | | | | |
| Executive functioning | | | **0.206** | **0.001** | **-0.104** | **0.002** | **-0.358** | **0.001** | **0.112** | **0.001** |

Coefficients in bold are significant after FDR correction (adopting a 5% false discovery rate) for 52 tests.
ADHD = attention-deficit hyperactivity disorder/sluggish cognitive tempo
Continuous dispositional and executive functioning measures were normalized to a mean of 0 and standard deviation of 1.

Table S13. Results of two sensitivity tests in which (a) all participants, including those with missing imputed cognitive test scores, or (b) only participants with no missing data after listwise deletion were used to define the latent executive functioning score which was regressed on simultaneously regressions the latent general factor and three specific factors defined in a bifactor model and on demographic covariates of no interest (age, sex, and race-ethnicity) in the second split half of the ABCD Study sample.

| Bifactor Model with Three Specific Factors | | | | | | | | |
|---|---|---|---|---|---|---|---|---|
| Criterion Variable | Factors | | | | | | | |
| | General | | Internalizing | | Externalizing | | ADHD | |
| | B | P < | β | P < | β | P < | β | P < |
| *Latent General Executive Functioning Test Scores* | | | | | | | | |
| Listwise deletion of missing cognitive test scores (N = 5555) | | | | | | | | |
| Executive functioning | -0.100 | 0.001 | 0.026 | 0.184 | -0.210 | 0.001 | -0.320 | 0.001 |
| Imputed missing cognitive test scores (N = 5934) | | | | | | | | |
| Executive functioning | -0.090 | 0.001 | 0.019 | 0.293 | -0.246 | 0.001 | -0.330 | 0.001 |

ADHD = attention-deficit hyperactivity disorder/sluggish cognitive tempo
Ns are slightly reduced because of sparse missing data on demographic covariates of no interest.

| Table S14. Pearson correlations (95% confidence intervals) among estimated factor scores *within* each confirmatory bifactor and second-order factor model in the second split half of the wave 1 ABCD Study sample (N = 5934) | | | | |
|---|---|---|---|---|
| Bifactor Models | | | | |
| Bifactor + 2 specific factors | Specific Externalizing | Specific Internalizing | | |
| General | 0.177 (0.152-0.202)*** | 0.148 (0.123-0.173)*** | | |
| Specific externalizing | | -0.144 (-0.169—0.119)*** | | |
| Bifactor + 3 specific factors | Specific externalizing | Specific internalizing | Specific ADHD | |
| General | 0.159 (0.134-0.184)*** | 0.124 (0.099-0.149)*** | 0.059 (0.034-.085)*** | |
| Specific externalizing | | -0.268 (-0.291--0.244)** | -0.022 (-0.047-0.00) | |
| Specific internalizing | | | -0.059 (-0.084--0.033)*** | |
| Bifactor + 4 specific factors | Specific externalizing | Specific internalizing | Specific ADHD | Specific somatization |
| General | 0.170 (0.14-0.19)*** | 0.134 (0.109-0.159)*** | 0.080 (0.54-0.105)*** | 0.035 (0.010-0.61)*** |
| Specific externalizing | | -0.330 (-0.353--0.308)*** | -0.028 (-0.53--0.002)* | -0.139 (-0.164- -0.114)*** |
| Specific internalizing | | | -0.100(-0.126--0.075)*** | 0.125 (0.100-0.150)*** |
| Specific ADHD | | | | -0.086 (-0.111- -0.060)*** |
| Second-Order Models | | | | |
| Second-Order (on 3 factors) | Externalizing | Internalizing | ADHD | |
| General | 0.959 (0.957-0.961)*** | 0.850 (0.843-0.857)*** | 0.959 (0.956-0.961)*** | |
| Externalizing | | 0.761 (0.750-0.771)*** | 0.861 (0.854-0.867)*** | |
| Internalizing | | | 0.758 (0.750-0.769)*** | |
| Second-Order (on 4 factors) | Externalizing | Internalizing | ADHD | Somatization |
| General | 0.948 (0.945-0.950)*** | 0.896 (0.891-0.901)*** | 0.959 (0.957-0.961)*** | 0.698 (0.684-0.711)*** |
| Externalizing | | 0.792 (0.782-0.801)*** | 0.858 (0.851-0.864)*** | 0.611 (0.595-0.627)*** |
| Internalizing | | | 0.804 (0.795-0.813)*** | 0.650 (0.636-0.665)*** |
| ADHD | | | | 0.618 (0.601-0.633)*** |

ADHD = attention-deficit hyperactivity disorder/neurodevelopmental
Hier = hierarchical model.
*p < .05; **p < .01; ***p < .001

| Table S15. Full set of correlations (95% confidence intervals) between estimated cognate factor scores *across* different confirmatory factor models (i.e., bifactor and second-order) in the second split half of the wave 1 ABCD Study sample (N = 5934) | | | | | |
|---|---|---|---|---|---|
| | Bifactor+2 | Bifactor+3 | Bifactor+4 | 2nd Order on 3 factors | 2nd Order on 4 factors |
| General factor scores | | | | | |
| 1-Factor | 0.918 (0.914-0.922) | 0.974 (0.972-0.975) | 0.979 (0.978-0.980) | 0.993 (0.993-0.994) | 0.995 (0.995-0.995) |
| Bifactor+2 | | 0.940 (0.938-0.943) | 0.953 (0.951-0.956) | 0.902 (0.897-0.907) | 0.920 (0.916-0.924) |
| Bifactor+3 | | | 0.991 (0.991-0.992) | 0.972 (0.971-0.974) | 0.966 (0.964-0.968) |
| Bifactor+4 | | | | 0.972 (0.970-0.973) | 0.977 (0.975-0.978) |
| 2nd Order on 3 | | | | | 0.994 (0.994-0.995) |
| Externalizing factor scores | | | | | |
| Bifactor+2 | | 0.786 (0.776-0.796) | 0.800 (0.790-0.809) | 0.625 (0.609-0.640) | 0.604 (0.588-0.620) |
| Bifactor+3 | | | 0.981 (0.980-0.982) | 0.418 (0.396-0.438) | 0.405 (0.383-0.426) |
| Bifactor+4 | | | | 0.465 (0.445-0.485) | 0.449 (0.428-0.469) |
| 2nd Order on 3 | | | | | 0.998 (0.998-0.998) |
| Internalizing factor scores | | | | | |
| Bifactor+2 | | 0.928 (0.924-0.932) | 0.541 (0.523-0.559) | 0.515 (0.496-0.534) | 0.351 (0.328-0.373) |
| Bifactor+3 | | | 0.765 (0.754-0.776) | 0.619 (0.603-0.634) | 0.495 (0.476-0.514) |
| Bifactor+4 | | | | 0.532 (0.513-0.550) | 0.564 (0.545-0.582) |
| 2nd Order on 3 | | | | | 0.952 (0.950-0.955) |
| ADHD factor scores | | | | | |
| Bifactor+3 | | | 0.988 (0.988-0.989) | 0.470 (0.450-0.490) | 0.459 (0.438-0.479) |
| Bifactor+4 | | | | 0.477 (0.457-0.496) | 0.467 (0.447-0.487) |
| 2nd Order on 3 | | | | | 0.995 (0.995-0.995) |
| Somatization factor scores | | | | | |
| Bifactor+4 | | | | | 0.747 (0.736-0.758) |

All correlations significant at $p < .001$

Supplemental Figure S1. Scree plots for parallel analysis with Glorfield correction for data from the first split half of the wave 1 ABCD Study sample and simulated data.

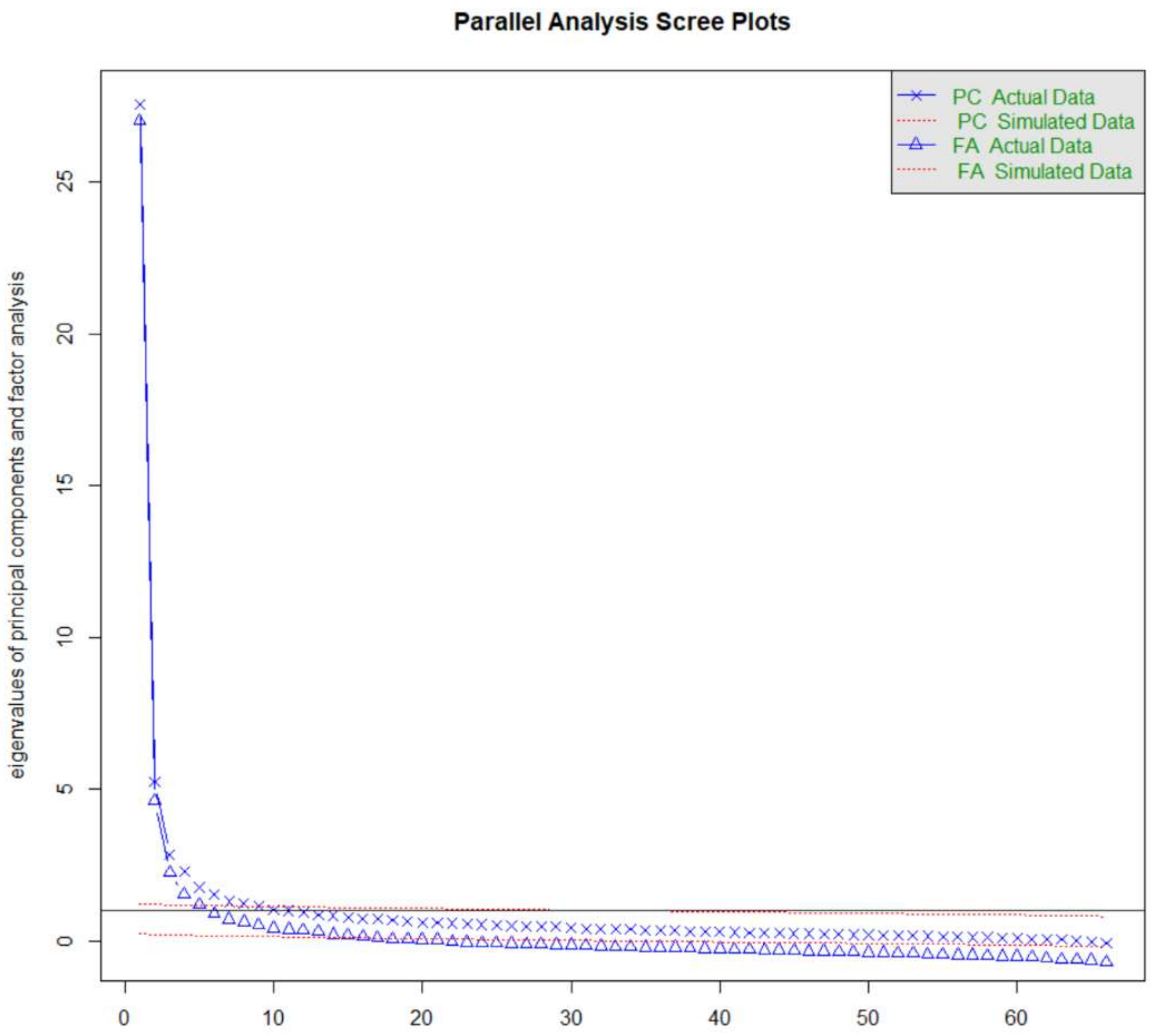

Supplemental Figure S2. Distributions of estimated general factors based on two different confirmatory models and differing numbers of lower-order factors (note differences in the scales of the x and y axes).

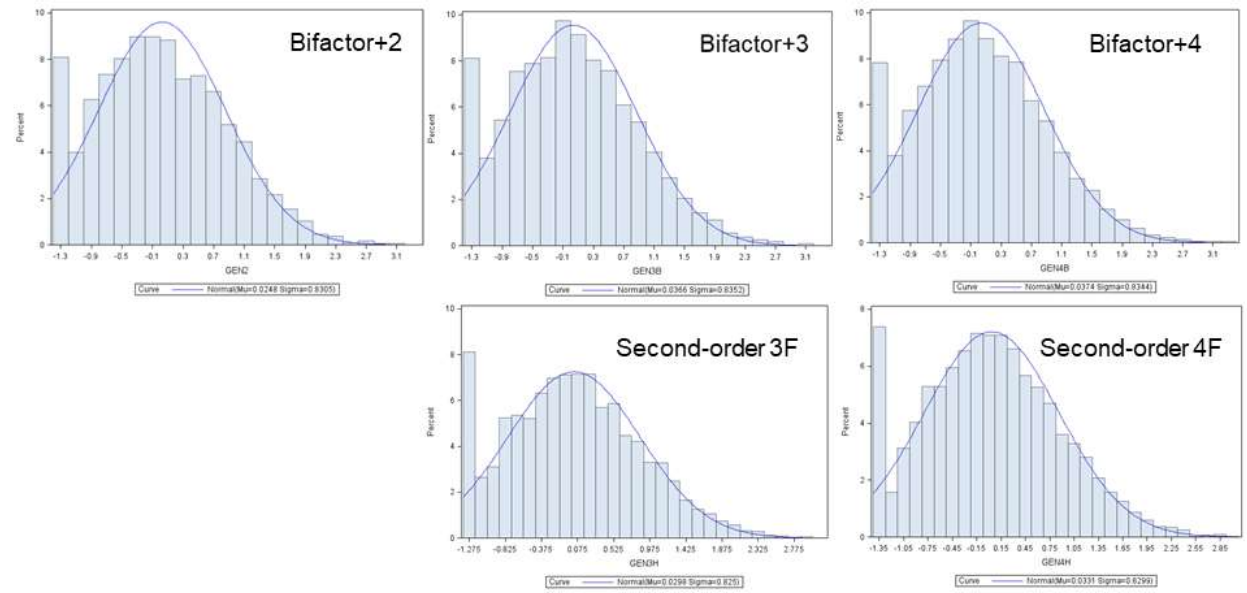

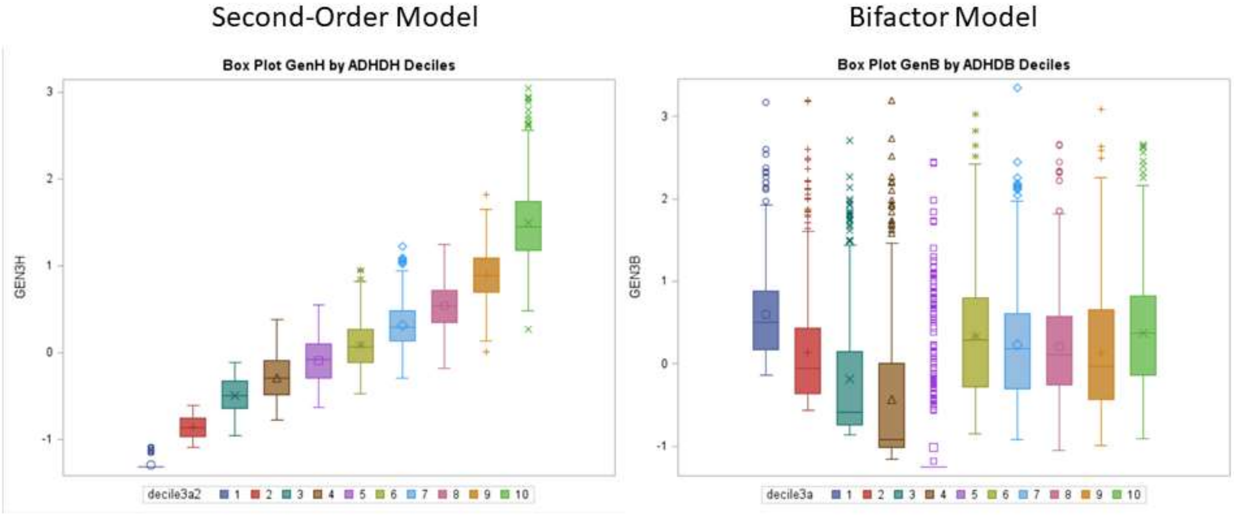

Supplemental Figure S3. Box plots of estimated general factor scores within deciles of estimated ADHD scores, separately for second-order and bifactor models.

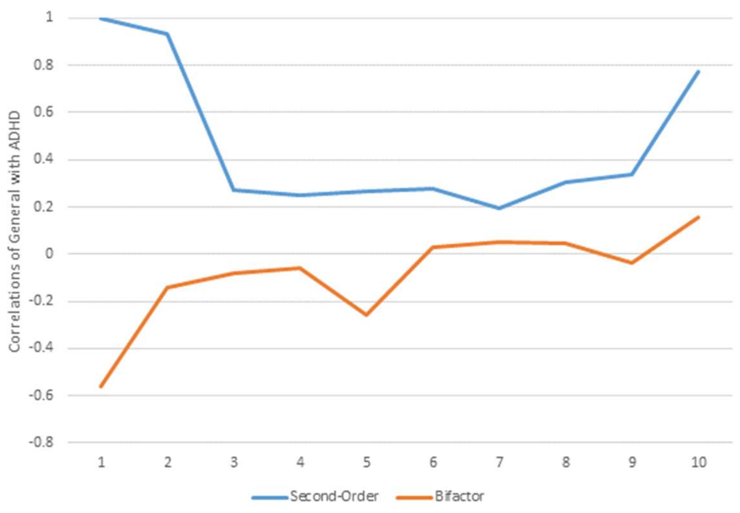

Supplemental Figure S4. Correlations within deciles of estimated ADHD factor scores with estimated general factor scores, separately for second-order and bifactor models.
